# Leaf- and diverged shoot meristem programs shape the stem in rice

**DOI:** 10.1101/2023.12.05.569800

**Authors:** Katsutoshi Tsuda, Akiteru Maeno, Ayako Otake, Kae Kato, Wakana Tanaka, Ken-Ichiro Hibara, Ken-Ichi Nonomura

## Abstract

The stem is the shoot axis in seed plants and has been a primary target in breeding to regulate crop height. However, early processes of stem development remain elusive. Here we show that regulators for shoot meristems and leaves determine the node-internode pattern in rice. Mutants of *KNOX1* genes *OSH15* and *OSH1*, known to maintain shoot meristem indeterminacy^1,2^, showed dwarfism due to enlarged nodes and diminished internodes. These genes confine node differentiation by repressing leaf developmental regulator *YABBY* genes in internodal vasculatures. *YABBY* expression, which normally extends from leaves to nodes along vasculatures, promotes nodal vascular differentiation and limits stem elongation. It expands in *knox1* mutants, and the loss of *YABBY* genes reverts their dwarfism. *OSH15* also represses node-specific *KNOX1* subclade genes *OSH6* and *OSH71* to allow internode elongation. Importantly, both *YABBY* and node-specific *KNOX1* genes are required for pulvinus formation at the leaf base, further elaborating the nodal structure for gravitropism. Thus, intersections between leaf and sub-functionalized shoot meristem programs shape nodes and internodes along the stem. Phylogenetic analysis showed that *KNOX1* sub-functionalization likely occurred in the progenitor of gymnosperms and angiosperms. Given that seed plants acquired their leaves independently from other vascular plant lineages^3,4^, the emergence of the node-internode pattern and *KNOX1*-*YABBY* regulatory module may be linked to seed plant leaf evolution.

## Introduction

Nodes and internodes are building units of the stem commonly found in seed plants. Nodes are attachment points of leaves to stems in which a complex vascular network allows the exchange of water and solutes. Internodes develop between nodes and greatly elongate to lift leaves and inflorescences for light capture and pollen dispersal, respectively. Regulation of internode elongation is important in crop breeding, as exemplified by semi-dwarf mutations utilized in the Green Revolution in the 1960s^5^. Despite their importance, however, stem development has been poorly studied compared to other organs such as leaves, flowers, and roots. This is probably due to the lack of clear landmarks in stem morphology in many species^6^. Grasses, including rice, are excellent models for studying stem development because distinctions among stem domains (nodes and internodes) and neighboring organs are clear^7,8^ (**Fig. 1a**).

**Figure 1.**
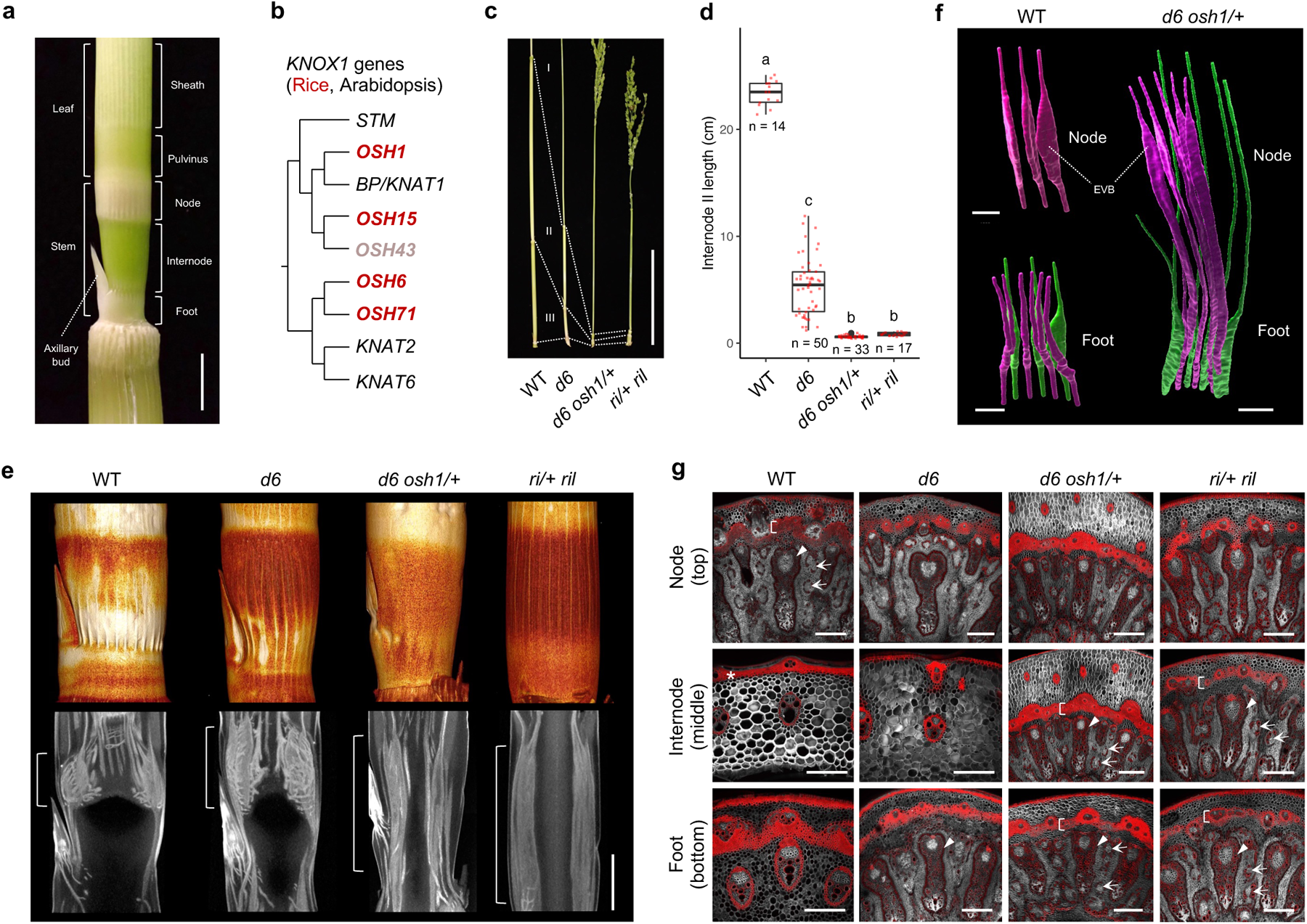
knox1 and blh mutants show enlarged nodes and diminished internodes. **a**, Structure of the young rice stem. **b**, The phylogenic relationship of rice (red) and Arabidopsis (black) *KNOX1* genes. *OSH43* is expressed at very low levels in the stem and excluded from the analysis. **c**, Shortened stem phenotypes of *knox1* and *blh* mutants. Roman numbers indicate internodes counted from the top. **d**, Internode II length at maturity. Different letters indicate significant differences (*p* < 0.05, Tukey-Kramer test). Red points, each sample; black points, outliers; center line, median; box limits, upper and lower quartiles; whiskers, 1.5x interquartile range. **e**, Micro-CT observation of young stems. The upper panels are external views (3D volume rendering images), and the lower panels are internal views of longitudinal sections. Brackets represent regions occupied by enlarged vascular bundles (EVBs). **f**, Vein structures extracted from micro-CT data. Veins in magenta are of the flag leaf phytomer, and those in green belong to one phytomer below. **g**, Transverse sections of mature stems. Arrows, arrowheads, and brackets indicate diffusing vascular bundles, EVBs, and lignified endodermal layers, respectively. An asterisk indicates a highly lignified epidermal layer in wild-type internodes, which was absent in all mutants. Confocal images for basic fuchsin (red) and calcofluor white (gray) were merged. Bars are 5 mm in **a**, 10 cm in **c**, 2 mm in **e**, 1 mm in **f**, and 200 µm in **g**.

In seed plants, the stem is produced from the shoot apical meristem (SAM) as a part of the phytomer, a developmental unit consisting of a leaf, a stem, and an axillary bud. In rice, we recently reported that cell fates for nodes and internodes are established stepwise^8^. At the flank of the shoot apical meristem (SAM), the cell fate for nodes starts being established, leading to the divergence of cell fates for leaves and stems. Subsequently, cells destined for internodes emerge from a limited number of cells. Thus, the cell fate for nodes is established earlier than the internodes, and the internode is the last domain whose cell fate is determined^8^. However, the molecular mechanisms underlying these processes are elusive.

Shoot systems with leaves evolved multiple times independently during the evolution of vascular plants. Fossil records indicate that leafless, branching sporophytes with terminal sporangia existed in all major lineages of vascular plants, including seed plants, ferns, and lycophytes^3,4^. Although leaves and leaf-like organs (i.e. leaves in seed plants, fronds in ferns, and microphyls in lycophytes) play common roles as lateral lamina for photosynthesis, their distinct ontogenies further support independent origins of these organs^3,4^. These facts suggest that the stem is derived from leafless shoot axes of ancestors and that the node-internode pattern emerged upon their leaf acquisition in seed plants.

Class I Knotted1-like homeobox (KNOX1) and BELL1-like homeobox (BLH) transcription factors (TFs) play important roles in the shoot axis in land plants^1,2^. In bryophytes, *KNOX1* genes promote sporophyte cell proliferation^9^. In vascular plants, their expression in shoot apical meristems and stem vasculatures is conserved^1,10–12^. Studies in angiosperms have revealed that *KNOX1* and *BLH* genes are essential for SAM maintenance and stem development^1,2,13–17^. Due to the striking phenotypes in mutants, their functions in the SAM have been extensively studied. However, their roles in stem development remain elusive.

In this study, we conducted genetic analyses to determine the function of *KNOX1* and *BLH* genes and explored key downstream targets in stem development. We uncovered the genetic basis of node and internode patterning with unexpected roles of regulators for leaf- and sub-functionalized shoot meristem regulators. Finally, a phylogenetic analysis for *KNOX1* genes provided an insight into stem evolution which may be associated with leaf acquisition in seed plants.

## Results

### *KNOX*1 and *BLH* genes restrict node differentiation

The rice stem consists of non-elongating and elongating domains (**Fig. 1a, Supplemental Fig 1a,b**). The node does not elongate but develops complex vascular networks between leaves and stems (**Supplemental Fig. 1c**). Internally, enlarged vascular bundles (EVBs), the node-specific vasculatures that are radially expanded, are especially conspicuous (**Fig.1** **f,g, Supplemental Fig. 1d**). EVBs have enlarged phloem surrounded by xylem, and are continuous to vasculatures of the leaf at the corresponding node. Diffusing vascular bundles (DVBs), that connect to veins from leaves two phytomers above in rice, surround and connect to EVBs (arrows in **Fig. 1g, Supplemental Fig. 1d**). These specialized vasculatures are important for water and solute exchange through various transporters^18^. The internode greatly elongates by cell division and elongation. Vascular bundles are narrow and simply vertical in internodes, probably optimized for rapid elongation (**Fig. 1g** and **Supplemental Fig. 1c**). There is another non-elongating part at the bottom, which we recently named the foot (**Fig. 1a**) ^7,8^. The foot also does not elongate but forms vascular networks connected to the axillary buds (**Supplemental Fig. 1b,c**).

We performed genetic analyses of two rice *KNOX1* genes *Oryza sativa homeobox15* (*OSH15*) and *OSH1*, which belong to the *BREVIPEDICELLUS* (*BP*) subclade (**Fig. 1b**). Loss-of-function mutants of *OSH15* (known as *d6* mutants) show dwarfism with abnormally shortened internodes, similar to Arabidopsis *bp* mutants (**Fig. 1c, d, Supplemental Fig. 1e,f**) ^16,17^. *osh1* mutants terminate the SAM after germination, and therefore, its function in stem development was unknown^14^.

Since *d6 osh1* double homozygotes are embryonic lethal^14^, we produced *d6 osh1*/+ double mutants to test whether *OSH1* is also involved in stem development. They showed an extremely short stem phenotype, indicating that *OSH1* is also important for stem development (**Fig. 1c, d, Supplemental Fig. 1e,f**). We also performed a genetic analysis of mutants of *BLH* cofactor genes *VERTICILLATE RACHIS* (*RI*) and *RI-Like* (*RIL*) ^19,20^. Their single and double mutants showed a range of dwarfism, and *ri*/+ *ril* double mutants developed extremely short stems similar to *d6 osh1*/+ (*ri ril* double homozygotes are also embryonic lethal^19^) (**Fig. 1c, d, Supplemental Fig. 1e,f**).

To understand the cause of these defects, we observed young stems using micro-computed tomography (micro-CT). Externally, *d6* single mutants had enlarged nodes and shortened internodes compared to wild types, and *d6 osh1*/+ and *ri*/+ *ril* lacked clear internodes (**Fig. 1e**). Internally, the node-specific EVBs were expanded in *d6* compared to the wild type, and ectopic EVBs were formed in the foot (**Fig. 1e,g**). Strikingly, EVBs occupied the entire stem in *d6 osh1*/+ and *ri*/+ *ril* double mutants, and tissue sections confirmed this occupancy by ectopic EVBs (**Fig. 1f,g**). In addition, less-lignified epidermis, highly lignified endodermal layers, and DVBs, which are features found in wild-type nodes, were observed all along the stem length in these double mutants (**Fig. 1g**). These mutant phenotypes suggest that KNOX1 and BLH TFs restrict the region that differentiates into nodes, thereby allowing internode formation.

### *YABBY* genes caused dwarfism in *knox1* and *blh* mutants

To unveil the molecular basis of node and internode patterning defects in these mutants, we dissected developing stems into transverse slices and compared transcriptome between the wild type and *d6* mutants (**Supplemental Fig.2a**). By clustering differentially expressed genes among tissues and genotypes, we identified two large gene clusters with distinct trends (**Supplemental Fig.2b**). In one cluster, genes were abundantly expressed in intercalary meristems (IM, a young internode) in the wild type and down-regulated in *d6* (cluster 3 in **Supplemental Fig.2b**). This cluster included many genes related to cell cycle and DNA synthesis as well as genes whose mutations are known to cause dwarfism^21–24^, suggesting that genetic pathways for internode formation were globally repressed in *d6* mutants (**Supplemental Figure 2c,d,e**). In another cluster, genes were highly expressed in the node and foot in the wild type, and their expression domains expanded toward IMs in *d6* mutants (cluster 2 in **Supplemental Fig.2b**). They included many cell wall- and lignin synthesis-related genes (**Supplemental Fig. 2c,d**). These trends were consistent with the *d6* phenotype with enlarged nodes and diminished internodes and with known *KNOX1* function to repress lignin pathways^25^.

To identify regulatory factors involved in the *d6* dwarfism, we explored TFs whose expression was altered in *d6*. Among TF genes, the *YABBY* family was exceptional because most members of this family were up-regulated in *d6* mutants (**Fig. 2a, Supplemental Fig 2c,f**). YABBY TFs are well-characterized regulators for lateral organ development^26–29^. A loss-of-function mutation in the Arabidopsis *YABBY* gene *FILAMENTOUS FLOWER* (*FIL*) is known to suppress the downward silique phenotype of *bp* mutants and slightly attenuate *bp* dwarfism in the *erecta* mutant background^30^, suggesting that there is a genetic interaction between *KNOX1* and *YABBY* genes. We knocked out five rice *YABBY* genes and found that mutations in two closely related *FIL*-like genes *TONGARI-BOUSHI1* (*TOB1*) and *TOB2*, which are important in spikelet development in rice^28,29^, greatly recovered internode growth in the *d6* mutant background, whereas mutations in other genes had minor effects (**Fig. 2b,c, Supplemental Fig. 3a)**. Therefore, we focused on *TOB1* and *TOB2* in subsequent analyses. Surprisingly, mutations both in *TOB1* and *TOB2* suppressed the extreme dwarfism both in *d6 osh1*/+ and *ri*/+ *ril* double mutants (**Fig. 2b,c, Supplemental Fig. 3a-c**). Histological observation confirmed that the internode formation was restored in *d6 osh1*/+ *tob1 tob2* quadruple mutants (**Fig. 2d**). Conversely, overexpression of *TOB1* or *TOB2* under the maize ubiquitin promoter resulted in extremely short stem phenotypes with expanded EVB formation (**Supplemental Fig. 3d-o**). These results suggested that ectopic expression of *TOB1* and *TOB2* was the major cause of dwarfism in *knox1* and *blh* mutants.

**Figure 2.**
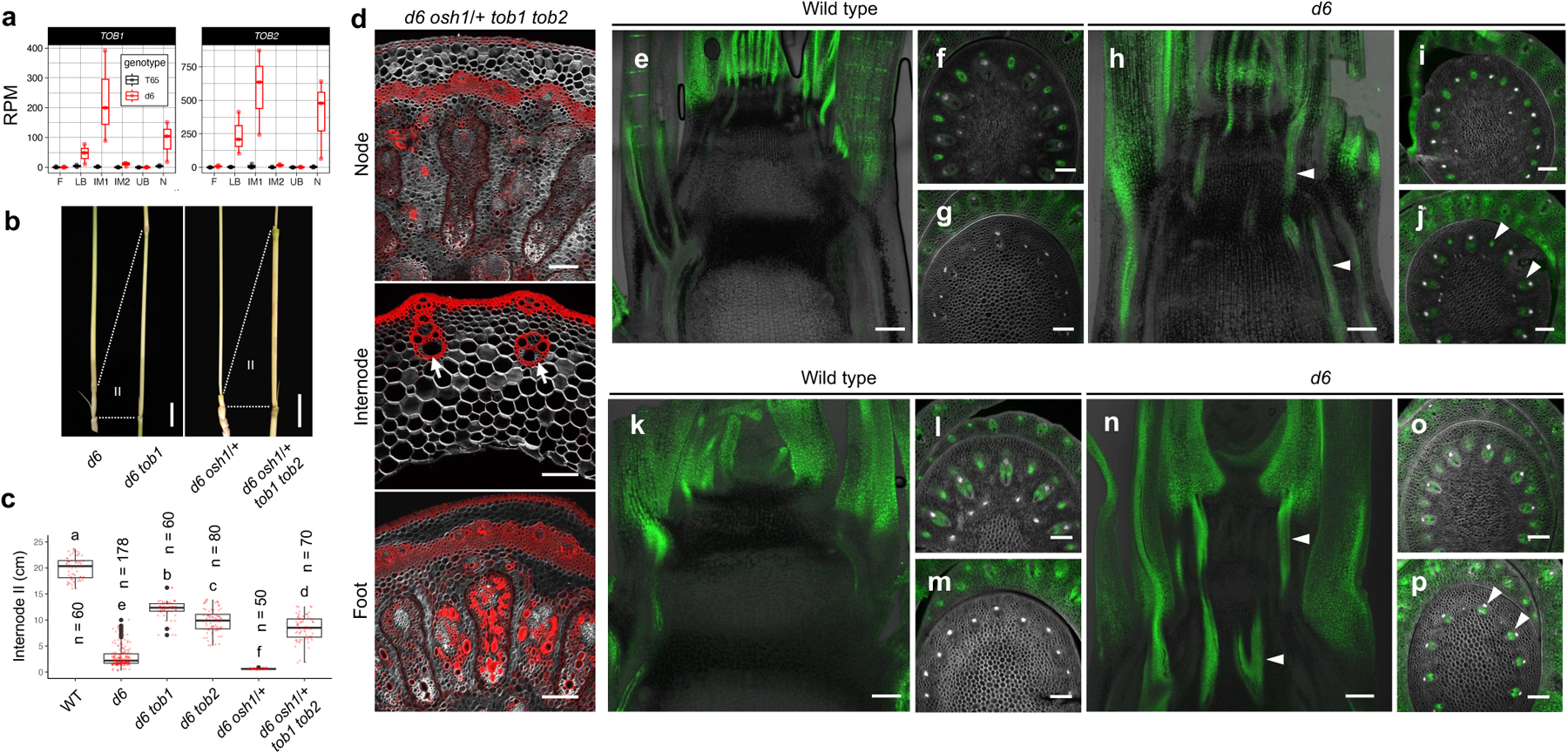
Ectopic expression of *YABBY* genes caused dwarfism in *knox1* mutants. **a**, Expression levels of *TOB1* and *TOB2* in mRNAseq data. Sample tissues are defined in **Supplemental** Fig. 2a. F, foot; LB, lower boundary; IM1, intercalary meristem1; IM2, intercalary meristem2; UB, upper boundary; N, node. Three biological replicates for each genotype. **b**, Mature internode II of *knox1* and *yabby* mutants. Dashed lines and Roman numbers represent internode II. **c**, Length of mature internode II in *knox1* and *yabby* mutants. Different letters indicate significant differences (*p* < 0.05, Tukey-Kramer test). **d**, Transverse sections of mature stems of *d6 osh1*/+ *tob1 tob2* quadruple mutants. Confocal images for basic fuchsin (red) and calcofluor white (gray) were merged. Arrows indicate normal vascular bundles restored in the quadruple mutant internodes. **e**-**p**, GFP fluorescence of *gTOB1-GFP* (**e**-**j**) and *gTOB2-GFP* (**k**-**p**) reporters. Background genotypes are shown above panels. **e**,**h**,**k**,**n**, Longitudinal sections of shoot apex at flowering transitions. Blight fields and GFP images were merged. **f**,**i**,**l**,**o**, Transverse sections of young nodes. **g**,**j**,**m**,**p**, Transverse sections of young internodes. Arrowheads indicate ectopic expression in internode vasculatures. Confocal images for calcofluor white (gray) and GFP (green) were merged. Bars are 2 cm in **b**, 200 µm in **d**, and 100 µm in **e**-**p**. In **a**,**c**, red and black transparent points, each sample; black points, outliers; center line, median; box limits, upper and lower quartiles; whiskers, 1.5x interquartile range.

Next, we investigated expression patterns of these *YABBY* and *KNOX1* genes in developing stems using genomic GFP reporters. In the wild type, *TOB1* and *TOB2* were broadly expressed in leaf primordia but not in shoot meristems and ground tissues of young stems (**Fig. 2e,k, Supplemental Fig. 4m**). Transverse sections revealed that both genes were broadly expressed beneath the youngest two leaf primordia (plastochron 1 (P1) and P2) (**Supplemental Fig 4.g,j**) and remained strongly expressed in provascular bundles (PBs) of nodes beneath P3 to P5, which will differentiate into EVBs (**Fig. 2f,l, Supplemental Fig. 4h,k**). In proximal regions corresponding to developing internodes, both *TOB1* and *TOB2* were repressed (**Fig. 2g,m, Supplemental Fig. 4i,l**). In contrast, *OSH15* and *OSH1* were broadly expressed in entire young stems including ground tissues and PBs (**Supplemental Fig. 4a-f**). One exception was their down-regulation in nodal PBs, where *TOB1* and *TOB2* were strongly expressed (**Supplemental Fig. 4b,e**). Observation of a *gTOB1-mCherry* and *gGFP-OSH15* double reporter line revealed their complementary accumulation patterns at nodes (**Supplemental Fig. 4m,n**). Thus, these *YABBY* and *KNOX1* genes are expressed in a mutually exclusive manner in the wild-type shoot apex.

In *d6* mutants, *TOB1* and *TOB2* were ectopically expressed along PBs: their expressions in PBs below the P3 to P5 leaf primordia were continuous from nodes to the bottom of the stem (**Fig. 2h-j, n-p**). Similarly, strong ectopic expressions of *TOB1* were observed in PBs of in young stems of *d6 osh1*/+ and *ri*/+ *ril* double mutants (**Supplemental Fig. 5**). Together, these observations showed that *YABBY* genes are expressed in nodal PBs and repressed by KNOX1 and BLH TFs in PBs of future internodes.

### Repression of *TOB1* is essential for stem development

Chromatin-immunoprecipitation followed by sequencing using *gGFP-OSH15* stem samples and anti-GFP antibodies suggested that OSH15 directly binds to *TOB1* and *TOB2* loci (**Fig.3a, Supplemental Fig. 6a**). Sequence comparisons around OSH15-bound regions in the *TOB1* promoter identified putative cis-regulatory elements evolutionarily conserved among grasses (**Fig. 3a, Supplemental Fig.6b**). Previously, we identified the consensus KNOX1 binding motif as two core GAs with three nucleotides apart (GAnnnGA) from OSH1 ChIPseq data^31^, and we found five such motifs highly conserved among grasses (**Supplemental Fig. 6b**). Electrophoretic mobility shift assays (EMSA) showed that OSH15-RI complex bound to these motifs, while either OSH15 or RI alone did not, indicating the requirement of both KNOX1 and BLH proteins to bind this motif (**Fig. 3b**). Mutations in these motifs abolished the binding. More importantly, when these conserved motifs were mutated in the *TOB1* genomic construct (*gTOB1-GFP_proMUT*, **Fig3a**), *TOB1-GFP* was ectopically expressed in PBs in developing internodes and caused severe dwarfism in the wild-type background (**Fig. 3c-g**). Transverse sections revealed that EVBs occupied the entire length of their stem at maturity (**Fig. 3h-j**). Collectively, the repression of *TOB1* by *OSH15* through evolutionarily conserved cis-elements is essential to restrict node differentiation.

**Figure 3.**
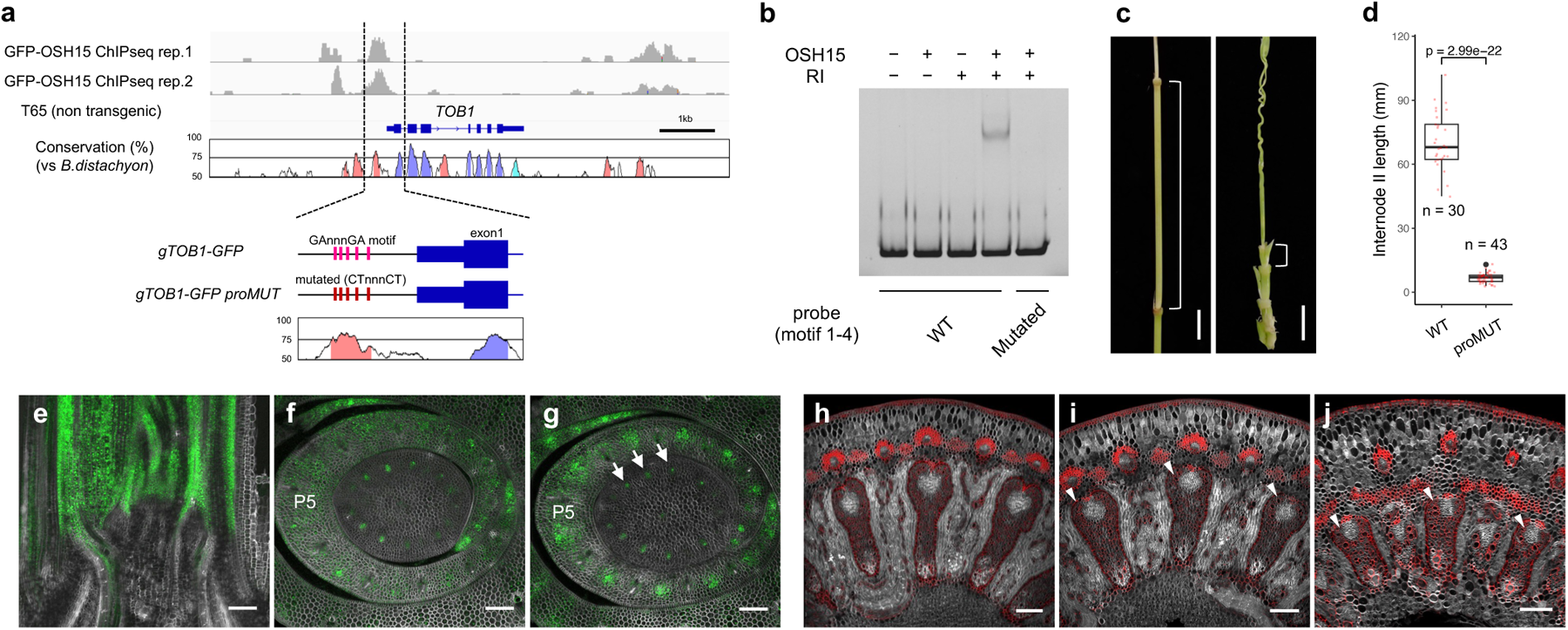
OSH15 directly represses *TOB1* to allow internode formation. **a,** OSH15-binding landscapes and nucleotide conservation at *TOB1* locus. ChIPseq data (two gGFP-OSH15 biological replicates and a T65 negative control) are shown above the gene model, and sequence conservation (vs Brachypodium distachyon) is shown below. Putative KNOX1 binding motif (GAnnnGA) and their mutations (CTnnnCT) introduced in the *gTOB1-GFP* construct are shown at the bottom. Regions in which the nucleotide conservation exceeded 75% are colored in pink, blue, and light blue for intergenic, coding, and UTR regions, respectively. **b**, EMSA assay using OSH15 and RI proteins and DNA fragments of the *TOB1* promoter containing putative KNOX-binding motif 1 to 4 (see **Supplemental Fig.6**). In the mutated probe, these conserved motifs were mutated into CTnnnCT. **c**, Mature stems of *gTOB1-GFP* (left) and *gTOB1-GFP_proMUT* (right) plants. Brackets indicate internode II. **d**, Length of internode II at maturity in transgenic plants shown in **c**. Red points, each sample; black points, outliers; center line, median; box limits, upper and lower quartiles; whiskers, 1.5x interquartile range. *p*-value represents a significant difference (two-sided t-test). **e**-**g**, GFP fluorescence in *gTOB1-GFP_proMUT* plants. **e**, longitudinal section; **f**, transverse section at P4 node; **g**, transverse section at P4 internode. Note that the periphery of the P4 internode is merging with P5 in **g**, indicating this section is at the bottom of the internode (compare with **Supplemental** Fig 4i). Arrows indicate ectopic expressions. Channels for calcofluor white (gray) and GFP (green) were merged. **h**-**j**, Transverse sections of *gTOB1-GFP_proMUT* mature stems corresponding to node I (**h**), internode II (**i**), and foot II (**j**). Channels for basic fuchsin (red) and calcofluor white (gray) were merged. Arrowheads are ectopically formed EVBs. Bars are 1 cm in **c**, 100 µm in **f**-**h**, 200 µm in **i**-**k**.

### Node-specific *KNOX1* genes also cause *d6* dwarfism

Results presented so far revealed the genetic interaction between *KNOX1* and *YABBY* genes during stem development. Previously, a similar genetic interaction was reported among *KNOX1* genes in Arabidopsis; the dwarfism of *bp* mutants was suppressed by mutations in *KNAT2* and *KNAT6*, members in the other subclade of *KNOX1*^32^. We found that *OSH6* and *OSH71*, two rice genes belonging to this clade (hereafter *KNAT2/6* clade), were upregulated in internodes of *d6* mutants (**Supplemental Fig. 7a,b**). As we expected, mutations in *OSH6* and *OSH71* suppressed *d6* dwarfism, showing that *OSH15* and *OSH6/71* genetically interact also in rice (**Fig. 4a**, **Supplemental Fig. 7c**). Genomic GFP reporters of OSH71 in wild-type backgrounds revealed that it accumulated in the SAM and in the entire young stem regions in the shoot apex, but are also expressed in the base of leaf primordia (**Fig. 4k**). In later stem development, OSH6 and OSH71 remained expressed in nodes but were down-regulated in internodes, leading to the node-specific expression pattern (**Fig. 4b-d**, **Supplemental Fig. 7d**). In *d6* mutants, both *OSH6* and *OSH71* were ectopically expressed in internodes (**Fig. 4e-g**, **Supplemental Fig. 7e**). Thus, *OSH6* and *OSH71* are node-specific *KNOX1* genes repressed by *OSH15*. These results are consistent with the report in *Arabidopsis*; *KNAT2* and *KNAT6* are expressed at nodes and are repressed by *BP* in internodes^32^.

**Figure 4.**
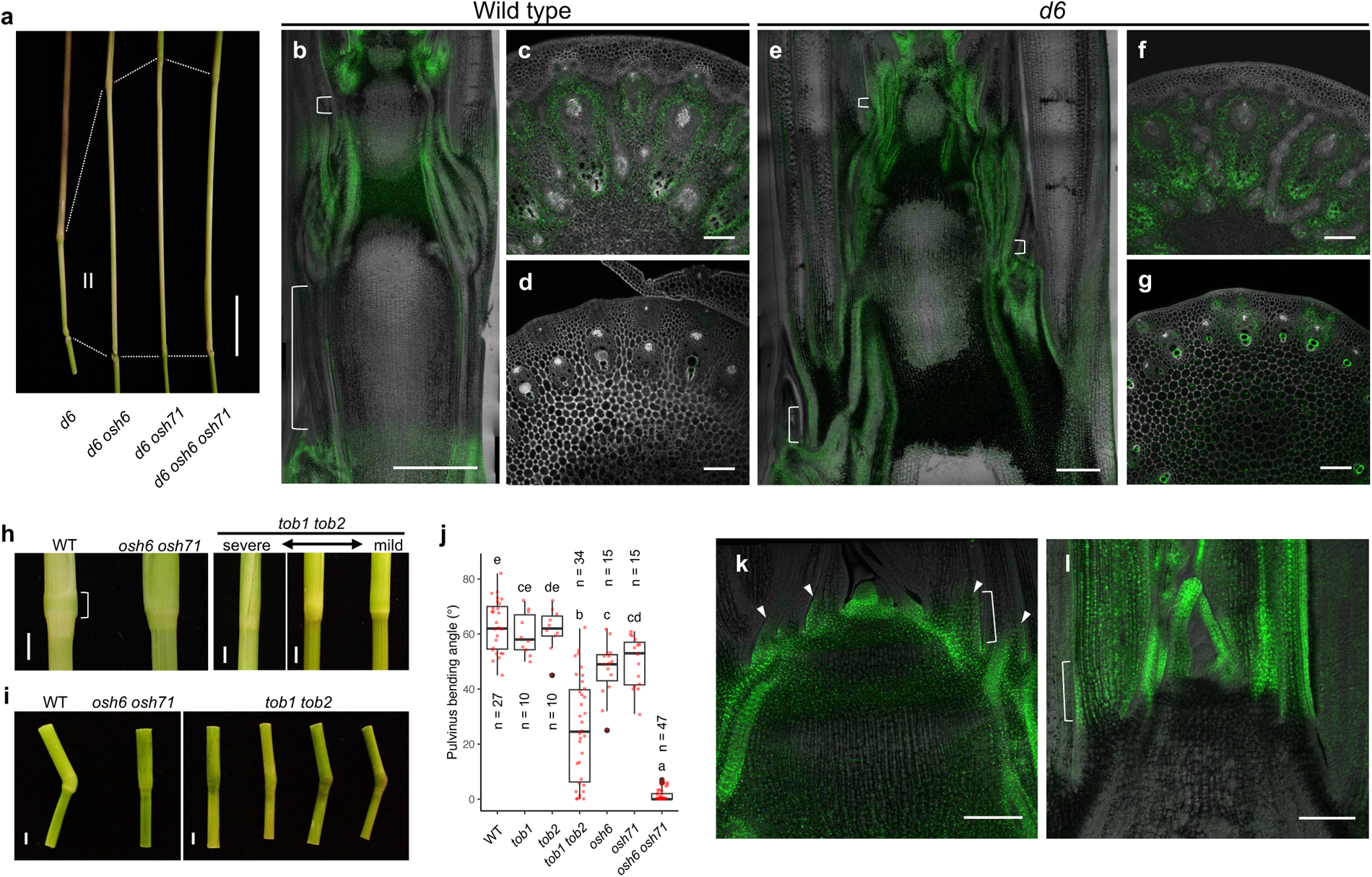
*OSH6* and *OSH71* are node-specific *KNOX1* genes essential for pulvinus development. **a**, Mutations in *OSH6* and *OSH71* attenuated dwarfism of *d6* mutants. Dashed lines and a Roman number indicate internode II. **b**-**g**, GFP fluorescence of *gGFP-OSH71* reporters in the wild type (**b**-**d**) and *d6* mutants (**e**-**g**). **b**,**e**, Longitudinal sections of developing stems. The blight field- and GFP channels were merged. **c**,**d,f**,**g**, Transverse sections of young nodes (**c**,**f**) and internodes (**d**,**g)**. Confocal images for calcofluor white (gray) and GFP (green) were merged. Brackets indicate developing internodes. **h**, Pulvinus-less phenotypes of *osh6 osh71* and *tob1 tob2* double mutants. **i**, Pulvinus bending tests. Samples were 10 day after the onset of gravity stimuli. **j**, Measurements of pulvinus bending angles after the gravity stimuli. Different letters indicate significant differences (*p* < 0.05, Tukey-Kramer test). Red points, each sample; black points, outliers; center line, median; box limits, upper and lower quartiles; whiskers, 1.5x interquartile range. **k**, *gGFP-OSH71* expression in the shoot apex. Arrowheads indicate *GFP-OSH71* accumulation at the base of leaf primordia. **l**, *gTOB1-GFP* expression in the shoot apex. Brackets in **k** and **j** are equivalent regions at the leaf base where expression domains of these genes overlap. The blight field and GFP channels were merged. Bars are 3 cm in **a**, 500 µm in **b**,**e**, 100 µm in **c**,**d**,**f**,**g**, 2 mm in **h**,**i**, and 200 µm in **k**,**l**.

### *OSH6/71* and *TOB1/2* are required for pulvinus formation

Interestingly, we noticed that *osh6 osh71* double mutants lacked the pulvinus, a characteristic structure at the leaf base adjacent to nodes (**Fig. 4h**). We recently reported that the pulvinus is the primary site exerting gravitropism in the rice stem^8^. *osh6 osh71* double mutants completely lost the pulvinus and the ability to bend nodes in response to gravity shift (**Fig. 4i,j**, **Supplemental Fig. 8a**). Surprisingly, we found that *tob1 tob2* double mutants also failed to form pulvini, although the phenotypic penetrance was 32.5 % (13 out of 40). *tob1 tob2* double mutant stems showed varying degrees of bending upon gravity stimuli; pulvinus-less plants could not bend the node, whereas milder plants did (**Fig. 4i,j**). In wild-type leaf sheaths, the epidermis and sclerenchymatous bundle caps of veins are highly lignified, and lysigenous aerenchyma forms between veins (**Supplemental Fig. 8b**). In wild-type pulvini, these aerenchymas were absent, lignification occurred only in xylem vessels, and collenchymatous bundle caps developed well. At leaf base in *osh6 osh71* and *tob1 tob2* double mutants with the pulvinus-less phenotype, all these characteristics of the pulvinus were absent, and tissue organization was nearly identical to that of the leaf sheath (**Supplemental Fig. 8b**). Thus, both *OSH6/71* and *TOB1/2* are important for pulvinus formation. Because expression domains of these genes partially overlap at the base of leaf primordia (**Fig. 4k,l**), the pulvinus is likely formed from where both regulators are co-expressed. Therefore, the pulvinus is a leaf domain whose development is governed by leaf and node regulators.

### *KNOX1* sub-functionalization occurred in early seed plants

Our data revealed that one of the *KNOX1* subclades and *YABBY* genes have functions in node development, and another *KNOX1* subclade represses their expression to allow internode formation. Importantly, *YABBY* genes are widely conserved only in seed plants ^33,34^, suggesting this family acquired prevalent functions in the ancestor of gymnosperms and angiosperms. We then questioned when the sub-functionalization of *KNOX1* genes occurred during plant evolution. We collected amino-acid sequences of KNOX1 proteins from land plant lineages and conducted a phylogenetic analysis (**Fig. 5**). In angiosperms, there are three major groups consisting of *STM*-, *BP*-, and *KNAT2/6*-subclades, each of which has functions partially overlapping but largely specialized for distinct domains along the shoot axis, i.e. shoot meristems, internodes, and nodes, respectively. All three *KNOX1* subclades are also present in gymnosperms but not in non-seed plants, suggesting that the sub-functionalization of *KNOX1* genes might have occurred in early seed plants and possibly linked to the acquisition of their leaves.

**Figure 5.**
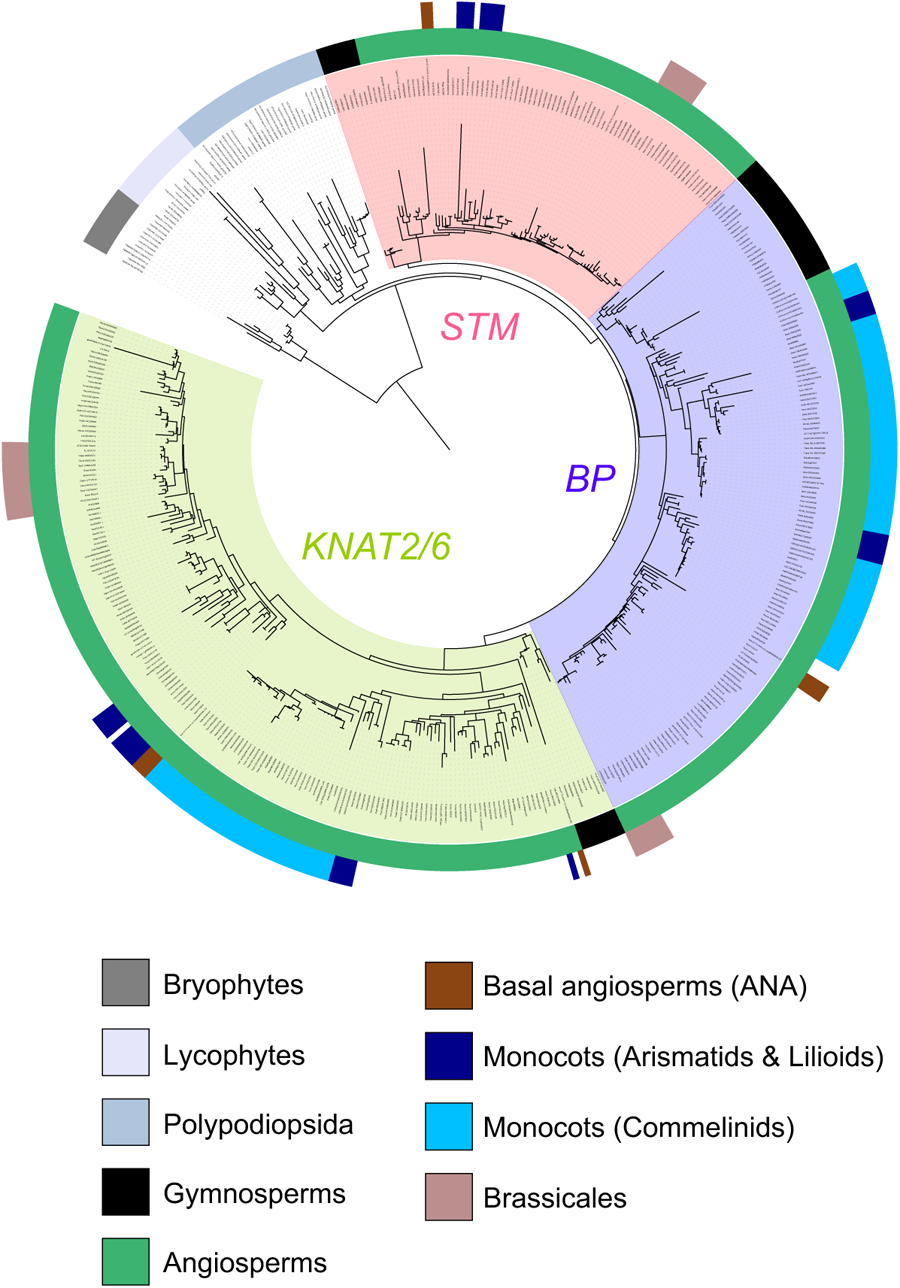
Three subclades of KNOX1 proteins of seed plants. A phylogenic tree of KNOX1 proteins from embryophytes. Shaded colors on the tree represent the *STM*, *BP*, and *KNAT2/6* subclades.

## Discussion

This study revealed a genetic mechanism that controls node-internode patterning in the stem (**Supplemental Fig. 9**). *YABBY* genes are broadly expressed at the boundary between leaf primordia and the young stem starting from one to two plastochrons after leaf initiation, but their expression becomes confined to future nodal PBs at later stages (**Supplemental Fig. 4g,h,j,k**). In proximal regions corresponding to future internodes, *YABBY* genes are repressed by the *BP*-clade *KNOX1* genes. This down-regulation is essential for internode development. Thus, the mutually exclusive pattern of *YABBY* and *BP* genes along the vascular bundle determines the node and internode pattern.

Although a range of evidence suggested that *TOB1* and *TOB2* induce EVB formation and limit internode elongation, it remains to be tested whether *YABBY* genes are required for these processes. We have observed neither extra stem elongation nor clear defects in EVBs in *tob1 tob2* double mutants so far. Further studies in combination with other *YABBY* mutations are required to establish their functions in nodal vascular differentiation.

Intriguingly, the ectopic expression of *YABBY* genes in *knox1* mutants was limited to the PBs of the internodes, but their developmental consequence included the entire internode. For example, tissue organization of nodes, including less lignified epidermal/subepidermal layers, highly lignified endodermis, and formation of DVBs, was observed throughout the longitudinal axis of *knox1* and *blh* mutant stems (**Fig.1g**). This suggests the existence of downstream targets of *YABBY* genes whose effect emanates from PBs to surrounding tissues non-cell autonomously. Besides, although vascular bundles in leaves also highly express these *YABBY* genes (**Supplemental Fig. 4g-l**), EVBs are limited to nodal vasculatures. Therefore, additional factor(s) present in developing stem should be necessary for EVB formation.

In this study, we focused on internode II, which is the last internode with well-developed foliage leaves. Interestingly, elongation of internode I subtending inflorescence was not largely affected in *knox1* and *blh* mutants (**Fig. 1c**). We found that *TOB1* and *TOB2* were down-regulated in the bract of inflorescence (**Supplemental Fig. 10**). This suggests that these *YABBY* genes are repressed by an unknown mechanism(s) in the bract and hence do not affect internode I.

In grasses in which nodes develop well, leaves encircle the entire circumference of the stem, and numerous vascular bundles directly enter the stem. In contrast, in most dicotyledons, leaves attach at a narrow point of the stem and fewer veins enter the stem. We speculate this is why the node is less conspicuous in many species. Nevertheless, the genetic interaction between *FIL* and *BP* in Arabidopsis^30^ and the dwarfism caused by dominant *FIL* mutations in cucurbits^35^ suggest that similar mechanisms for stem development operate in a wide range of species.

Besides the sub-functionalization of *KNOX1* genes, our phylogenetic analysis revealed interesting trajectories of their evolution (**Fig. 5**). *STM*, which is essential for SAM maintenance, was lost during monocot evolution; *STM* homologs were present in Alismatids and Lilioids (two paraphyletic groups in monocots) but were absent in Commelinids (a monophyletic group including Arecales, Poales, Zingiberales and Commelinales). A caveat is that genes from Arecales were, unfortunately, absent in our dataset. Previously, it was reported that *STM*-like genes were found in Asparagales and Arecales but not in Poales, including grasses ^36,37^. Therefore, the *STM* subclade was lost after the split of Arecales and other sister groups in Commelinids. Instead, *BP* genes duplicated at least once during monocotyledon evolution; one subset, including *OSH1* in rice and *kn1* in maize, had been deployed for SAM maintenance^14,15^, while another including *OSH15* might have kept its original function for internodes^17^. In the *KNAT2/6* subclade, there are two large branches and lineage-specific gene loss was again observed. For example, Commelinids lost one branch, while Brassicales, including Arabidopsis, lost another (**Fig. 5**). Therefore, functions between *KNAT2/6* subclade genes are not necessarily the same in Arabidopsis and in rice, although their genetic interactions with *BP* subclade genes are conserved.

Dwarf mutations in the gibberellin pathway have been utilized in the Green Revolution to achieve lodging resistance and yield gains for a long time. However, there are shortcomings such as low fertility under cool temperatures, narrow stem diameters, and overuse of chemical fertilizers leading to environmental burdens^5,38–40^. Therefore, alternative strategies to regulate crop height are important. It was recently reported that a dominant allele of a *YABBY* gene with an increased translation also resulted in semi-dwarfism in cucurbits^35^. Our data showed that cis-regulatory mutations in *YABBY* genes can affect stem length. It will be interesting to seek optimal allele(s) that achieve a desired height. Further efforts to understand the genetic basis of stem development and fine-tune its regulators would allow us to manipulate crop height for agricultural improvement.

## Methods

### Plant materials and growth conditions

For field-grown plants, a japonica rice variety Taichung65 (T65) was used as the wild type. The *d6* allele with *Tos17* retrotransposon insertion in the fourth exon of *OSH15* was used^17^. *osh1*, *ri*, and *ril*, mutants were described previously^14,19^. The original *tob1* allele^29^ was used for *d6 tob1* and *d6 osh1*/+ *tob1* mutant observation. Combinations of these mutants were generated by crossing and were genotyped using primers and restriction enzymes listed in **Supplemental Table1**. Plants were grown in the paddy field in Mishima, Shizuoka, Japan, in the summer of 2017-2022. Other combinations of mutations were generated using genome editing. Transgenic plants were grown in a closed greenhouse under natural light supplemented with a hydrogen lamp for 12 hours during the day. Nipponbare was used as the material and the wild-type control for transgenic studies. Temperature was kept at 32 °C during the day and at 25 °C at night. Relative humidity was maintained at 70%.

### Genome editing and rice transformation

Genome editing was performed using the system reported previously^41^. Agrobacterium-mediated rice transformation was conducted as described previously^42^. Japonica rice variety Nipponbare (Nip) was used as material and the wild type. Genomic DNA was extracted from regenerated T0 plants, target regions were PCR amplified, and mutations were identified by Sanger sequencing. Guide RNAs and induced mutations are listed in **Supplemental Table2**. We selected at least two independent transgenic lines for each construct and genetic background and observed phenotypes for at least two generations.

### Fluorescent reporter construction

The genomic fluorescent reporters for *OSH15* and *OSH1* were reported previously^14,43^ and other reporters were constructed in the same way. Briefly, BAC clones were digested with restriction enzymes, separated in 0.8% gel by electrophoresis, and genomic DNA fragments containing entire genes of interest with upstream and downstream regions were extracted from gel slices (enzymes and band sizes were listed in **Supplemental Table3**). pENTR2.2^43^ with overhangs matching the ends of genomic DNA fragments were amplified by inverse PCR, and the genomic DNA fragments from BAC clones were cloned using NEBuilder (NEB). Fluorescent protein-coding sequences were inserted in-frame at N- or C-terminus using inverse PCR and NEBuilder. The resultant genomic fragments with fluorescent fusions were transferred to binary vectors pPG or pPGn^43^ for hygromycin or G418 selection, respectively) using LR clonase II (Thermo Fisher). Reporter constructs were introduced into rice calli through Agrobacterium-mediated transformation^42^. Nucleotide sequences amplified by PCR were checked by Sanger sequencing. Primers were listed in **Supplemental Table1**.

### *TOB1* and *TOB2* overexpression

Full-length coding regions of *TOB1* and *TOB2* without stop codons were cloned into pENTR (Thermo Fisher) and transferred to pPUG2 for overexpression of C-terminal GFP fusion proteins^43^. Constructs were introduced into Nipponbare calli by Agrobacterium-mediated transformation^42^. We selected at least five independent lines that were GFP-positive in root apices and observed their phenotypes.

### *TOB1* promoter mutagenesis

*TOB1* promoter region (2.2 kb) was PCR amplified and cloned into pCR_Blunt vector (Thermo Fisher). This plasmid was used as a template to introduce mutations into putative KNOX1 binding site BS1 to BS3 by inverse PCR. The inverse PCR products were self-ligated and introduced into *E. coli*. After confirming the sequence, this plasmid was used as a template for the second inverse PCR to introduce additional mutations into BS4 and BS5. This PCR product was again self-ligated, and the sequence was confirmed. The resultant mutagenized promoter was introduced into the pENTR2.2_gTOB1-GFP in place of the original wild-type promoter. The resultant construct with promoter mutations was designated as gTOB1-GFP_proMUT. Primers were listed in **Supplemental Table1**. We selected seven independent gTOB1-GFP_proMUT lines that were GFP-positive and measured their internode length.

### Statistical analysis

For multiple comparisons, the Tukey-Kramer test was conducted. For paired comparison, the two-sided Student’s T test was performed. These analyses were performed in R 4.2.1.

### Measurements of the internode length

The length of internode II, which belongs to the flag leaf phytomer in rice, was measured at maturity when plants were at the seed ripening stage after flowering. The top three to four culms from at least four plants were chosen for each genotype. For *TOB1* and *TOB2* overexpressors and *gTOB1-GFP_proMUT* plants, that did not emerge inflorescences from the flag leaf, we removed leaves and measured the length after confirming no further leaf was being formed. Source data were listed in **Source Data1**.

### Stem bending assay

Plants at the seed ripening stage were laid down to the ground (90-degree rotation). After 10 days, stem bending angles at nodes I, II and III were recorded for each culm. The largest angles in each culm were chosen for plotting. Data were collected from at least 3 culms and 3 plants. Source data were listed in **Source Data1**.

### Histological observation

Stem samples at maturity were harvested and fixed in FAA (formalin: acetic acid: ethanol: water = 10:5:50:35) at 4 °C overnight, rehydrated in water, and cleared using ClearSee (Fujifilm) for more than two weeks. Upon observation, samples were water-washed and sectioned manually using razor blades. Sections were stained with 0.1% calcofluor white and 0.1% basic fuchsin for 15 min, briefly rinsed in water for 5 min, and mounted in 0.5 x water-diluted VECTASHIELD (Vector laboratories, H-1000). Confocal images were captured using Fluoview FV300 CLSM system (Olympus) with excitation at 404 nm and detection at 430-460 nm for calcofluor white and with excitation at 534.5 nm and detection at 585-640 nm for basic fuchsin. Because hand sections were not always flat and did not fit in a single focal plane, approximately 50 µm thick z-stack images were obtained at 5 µm intervals. Multiple z-stack images overlapping each other (at least 1/3 along the x- and y-axes) were captured for larger samples such as nodes, and stitched using the plug-in tool for pairwise stitching in Fiji 2.9.0^44^. For *TOB2* overexpressor observation, fresh samples were manually sectioned using razer blades, mounted in water, and directly observed under BX50 light microscope (Olympus) equipped with DP22 digital camera (Olympus).

### Fluorescent reporter observation

Shoot apices of reporter lines were dissected and fixed in 4% paraformaldehyde for 1 hour and embedded in 7% ultra low range agarose gel (BioRad). Fifty µm sections were prepared using a vibratome DTK-1000 (Dosaka) and mounted in 0.5 x water-diluted VECTASHIELD (supplemented with 0.01% calcofluor) and imaged using Fluoview FV300 CLSM system (Olympus). The excitation and detection wavelengths were 404 nm and 430-460 nm for calcofluor white, 488 nm and 505-525 nm for GFP, and 543.4 nm and 585-640 nm for basic fuchsin and mCherry. When the area of interest exceeded a single microscopic field of view, multiple overlapping images (at least 1/3 along the x- and y-axis) were taken and stitched using the plug-in tool for pairwise stitching in Fiji 2.9.0^44^. We observed at least three independent transgenic lines in each genetic background and confirmed that expression patterns were consistent among the same genotype.

### Micro-computed tomography

Micro-computed tomography (micro-CT) scanning was performed as described previously^20,45^. Rice stem samples (5 mm) were fixed in FAA (formalin:acetic acid: 50% [v/v] ethanol = 5:5:90) overnight. The fixative was replaced with 70% ethanol and stored at 4 °C until observation. Before scanning, samples were soaked in the contrast agent (1.0% [w/v] phosphotungstic acid in 70% [v/v] ethanol) for 6-11 days and scanned using X-ray micro-CT at a tube voltage peak of 85 kVp and a tube current of 90 μA. Samples were rotated 360° in steps of 0.24°-0.3°, generating 1,200-1,500 projection images of 992 × 992 pixels. The micro-CT data were reconstructed at an isotropic resolution of 5.8-9.7 μm^3^. Three-dimensional tomographic images were obtained using the OsiriX software. To visualize vascular networks, we harvested plants from the paddy field and immediately let plants absorb an iohexol containing contrasting reagent (Omnipaque, GEHealthcare) from the cut for 2 hour at room temperature. Samples were scanned using X-ray micro-CT at a tube voltage peak of 90 kVp and a tube current of 80 μA. Samples were rotated 360° in steps of 0.2°, generating 1,800 projection images of 992 × 992 pixels. The micro-CT data were reconstructed at an isotropic resolution of 9.0-9.5 μm^3^. Vascular networks were visualized by segmentation using Imaris software.

### Transcriptome analysis

Stem samples at the 10 mm stages were cut into 1-2 mm slices, and total RNAs were extracted using Nucleospin RNA Plant (Macherey-Nagel). Five hundred ng of total RNAs were used for each sample. mRNAseq libraries were constructed using KAPA RNA HyperPrep Kit (KAPA BIOSYSTEMS). Single-ended 100 bp libraries were sequenced using Hiseq4000. Read data were trimmed using trimmomatic-0.38^46^ with the following options: ILLUMINACLIP:TruSeq3-SE.fa:2:30:10 LEADING:3 TRAILING:3 SLIDINGWINDOW:4:15 MINLEN:30. Quality-filtered reads were mapped to MSU7.0 rice genome using hisat2^47^ with the default setting, and output bam files were converted to read count data using featureCounts^48^ implemented in Rsubread 1.28.1^49^. Genes with 10 reads or more, at least in 3 samples, were selected as expressed genes (25,350 genes). To extract genes whose expression levels dynamically changed among tissues and genotypes, differential gene expression analysis was performed for expressed genes using DEseq2^50^, and we selected genes with the adjusted *p*-value < 0.01, yielding 7,111 genes. These dynamically expressed genes were clustered using hclust function in base R (4.2.1) with k=7. For up-regulated and down-regulated genes in *d6* mutants, we extracted differentially expressed genes with |log2(fold change)| > 3. Functional category enrichment analysis was performed using GOseq^51^ with MapMan annotation data for Oryza sativa (Osa_MSU_v7) ^52^.

### Reverse transcriptase quantitative-PCR (RT-qPCR)

Five hundred ng of total RNAs were treated with DNase I and used for cDNA synthesis using Superscript III (Thermo Fisher) in 20 µL reactions. After synthesis, cDNAs were diluted with 80 µL of TE buffer. For quantitative RT-PCR, 1 µL of cDNA was used as a template. Quantitative PCR (qPCR) was performed in 10 µL reactions using KAPA SYBR Fast qPCR Kit (KAPA BIOSYSTEMS). Expression levels were calculated using the 2–ΔΔCt method. *Rice actin1* (*RAc1*) gene was used as an internal standard. Primers used in qPCR are listed in **Supplemental Tabel1**.

### Chromatin immunoprecipitation (ChIP) assay followed by sequencing and qPCR

ChIP assay was performed as described previously^14,31^ with the following modifications. Young stem samples (< 5 mm length) from *gGFP-OSH15* were cut vertically in half and fixed in 1% formaldehyde for 10 min at room temperature. Fixed samples were washed with distilled water three times and stored at −80 °C until the assay. Chromatin was extracted from at least 10 stem samples and 20 µg of chromatin was used per replicate. Immunoprecipitation was done using 1 µg of anti-GFP antibody (MBL) bound to Dynabeads protein A (Thermo Fisher). After washing and elution, qPCR was performed as described previously^14,31^ using KAPA SYBR Fast qPCR Kit and primers listed in **Supplemental Table 1**. For ChIP-seq study, paired-ended libraries were constructed using SMART ChIPseq kit (TAKARA) and sequenced using HiseqX. Read data were trimmed using fastp^53^ with the following options; -3 -f 3 -h -q 15 -n 10 -t 1 -T 1 -l 20. The trimmed reads were mapped using bwa mem^54^ with the default setting and visualized using IGV software^55^.

### *TOB1* promoter sequence analysis

Nucleotide sequence conservation around the *TOB1* locus was analyzed and visualized using VISTA-Point^56^ (https://genome.lbl.gov/vista/index.shtml). *TOB1* promoter sequences of *Brachypodium distachyon*, *Sorghum bicolor*, and *Setaria viridis* were retrieved from the VISTA-Point browser. For *Zea mays*, promoters of duplicated *TOB1* orthologous genes were searched using BLAST in Phytozome 13^57^ (https://phytozome-next.jgi.doe.gov/), and the top two hits were retrieved. Sequences were aligned with the muscle option and visualized using SeaView^58^.

### Electrophoretic mobility shift assay (EMSA)

Coding regions of *OSH15* and *RI* were PCR amplified and cloned into the pTnT vector (Promega). The FLAG and Myc tag were introduced into the N-termini of OSH15 and RI, respectively, for checking protein production. Proteins were produced using 5 µg of plasmids in 25 µL in vitro translation reaction of TnT SP6 High-Yield Wheat Germ Protein Expression System (Promega). For the OSH15-RI complex, 2.5 µg of each plasmid was used. Probes sequences were amplified from gTOB1-GFP and gTOB1-GFP_proMUT constructs and cloned into pCR_Blunt vectors (Thermo Fisher). Probes were synthesized by PCR using 5’-FAM labelled primers annealing to the vector (**Supplemental Table1**). The binding reactions were performed on ice for 30 min in a binding buffer (10 mM Tris-HCl pH7.5, 50 mM NaCl, 1% CHAPS, 0.5 mM EDTA, 1 mM MgCl_2_, 0.5 mM DTT, 60 µg/mL Salmon Sperm DNA, 8% glycerol) with 10 nM probes and 2 µL of in vitro translated proteins in 20 µL total volumes. Samples were loaded in 5% polyacrylamide gels, and electrophoresis was performed in 0.5 x TBE buffer at 100 V for 2 hrs at 4 °C. FAM-labelled probes were visualized using WSE-5600 CyanoView (ATTO).

### Phylogenetic analysis

Amino acid sequences of KNOX1 proteins were obtained from Phytozome 13 through BLAST search using the OSH1 sequence as a query against embryophyte databases. In addition, the same search was done in Plaza Gymnosperm database^59^. Hits with the following criterion were retained as KNOX1 proteins: more than 200 amino acids alignments (removing KNAT-M class without homeodomains), and smaller p-values than Arabidopsis KNAT3, KNAT4, and KNAT5 (removing distantly related class II KNOX). In total, 453 KNOX1 protein sequences were collected (**Source Data2**). These sequences were aligned using mafft online server (https://mafft.cbrc.jp/alignment/server/index.html) ^60^ using the option GINSi, and the multiple sequence alignment (MSA) was trimmed using TrimAl^61^ with -gt ranging from 0.6 to 0.95. ML trees were inferred using the iqtree web server (http://iqtree.cibiv.univie.ac.at/)^62^ with the default setting. The tree was visualized in iTOL (https://itol.embl.de/)^63^.

### Data availability

Transcriptome and ChIP-seq data are available in the DDBJ Sequenced Read Archive (DRA) under the accession numbers PRJDB13390 and PRJDB16754, respectively. Accession numbers of rice genes studied here are as follows: OSH1 (LOC_Os03g51690), OSH6 (LOC_Os01g19694), OSH15 (LOC_Os07g03770), OSH71 (LOC_Os05g03884), TOB1 (LOC_Os04g45330), TOB2 (LOC_Os02g42950), TOB3 (LOC_Os10g36420), YAB1 (LOC_Os07g06620), YAB2 (LOC_Os03g44710), YAB6 (LOC_Os12g42610).

## Acknowledgment

We thank Misako Kashihara for field management, Sarah Hake for thoughtful comments on the manuscript, Masaki Endo for providing genome editing vectors, and Hiroyuki Hirano for providing *tob1*, *ri*, and *ril* seeds. Computations were partially performed on the NIG supercomputer at ROIS National Institute of Genetics.

## Author contributions

K.T. designed this work, performed experiments, and analyzed data. A.M. conducted micro-CT observation. A.O., K.K., and K-I. H. contributed to experiments. W.T. constructed genome editing plasmids for *TOB* genes. K-I.N. supervised the project. K.T. wrote the manuscript with feedback from all authors.

## Materials & Correspondence

Correspondence and requests for materials should be addressed to K.T.

