## Supplemental Figure 1 to 10 for "Leaf- and diverged shoot meristem programs shape the stem in rice"

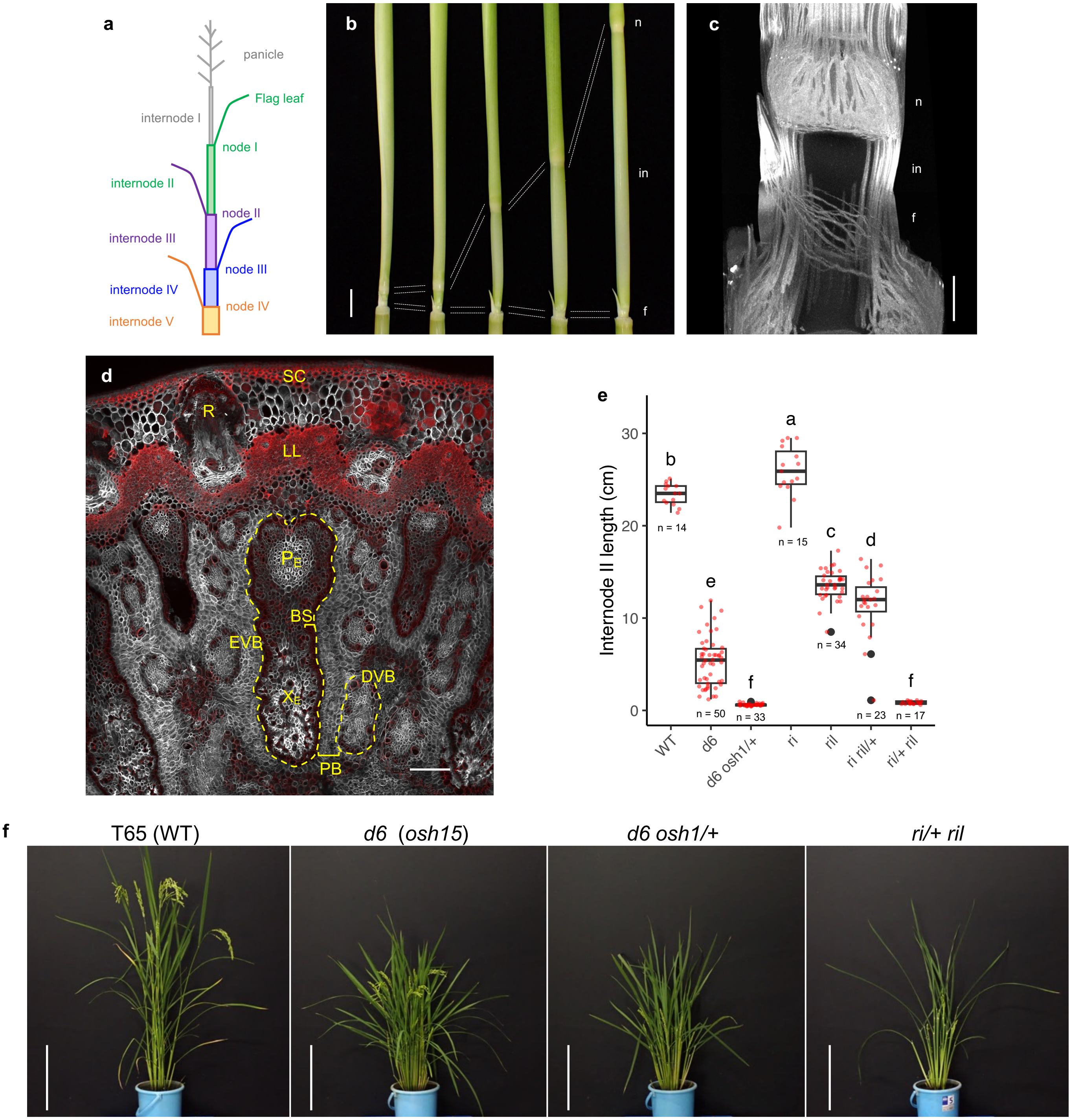


**Supplemental Figure 1. Structure of the rice stem and *knox1* and *blh* mutant phenotypes.** **a**, The schematic representation of rice plant structure and the numbering of each stem domain. Organs and stem domains belonging to the same phytomer have the same colors. **b**, Rice stems at various stages of elongation. **c**, A longitudinal view of stem inner structures imaged using micro-CT. Vascular networks are visible as X-ray-absorbing structures. **d**, A transverse section of the wild-type mature node. EVB and DVB are shown with dashed lines. Confocal images for basic fuchsin (red) and calcofluor white (gray) were merged. **e**, Length of internode II in *knox1* and *blh* mutants at maturity. Different letters indicate significant differences ($p < 0.05$, Tukey-Kramer test). Red points, each sample; black points, outliers; center line, median; box limits, upper and lower quartiles; whiskers, 1.5x interquartile range. **f**, Mature plants of *knox1* and *blh* mutants. n, node II; in, internode II; f, foot II; SC, sclerenchyma; R, crown root; LL, lignified layer (derived from endodermis); EVB, enlarged vascular bundle; $P_E$, Phloem of EVB; $X_E$, xylem of EVB; BS, bundle sheath; PB, parenchymal bridge; DVB, diffusing vascular bundle. Bars are 1 cm in **b**, 1 mm in **c**, 100 µm in **d**, and 30 cm in **f**.

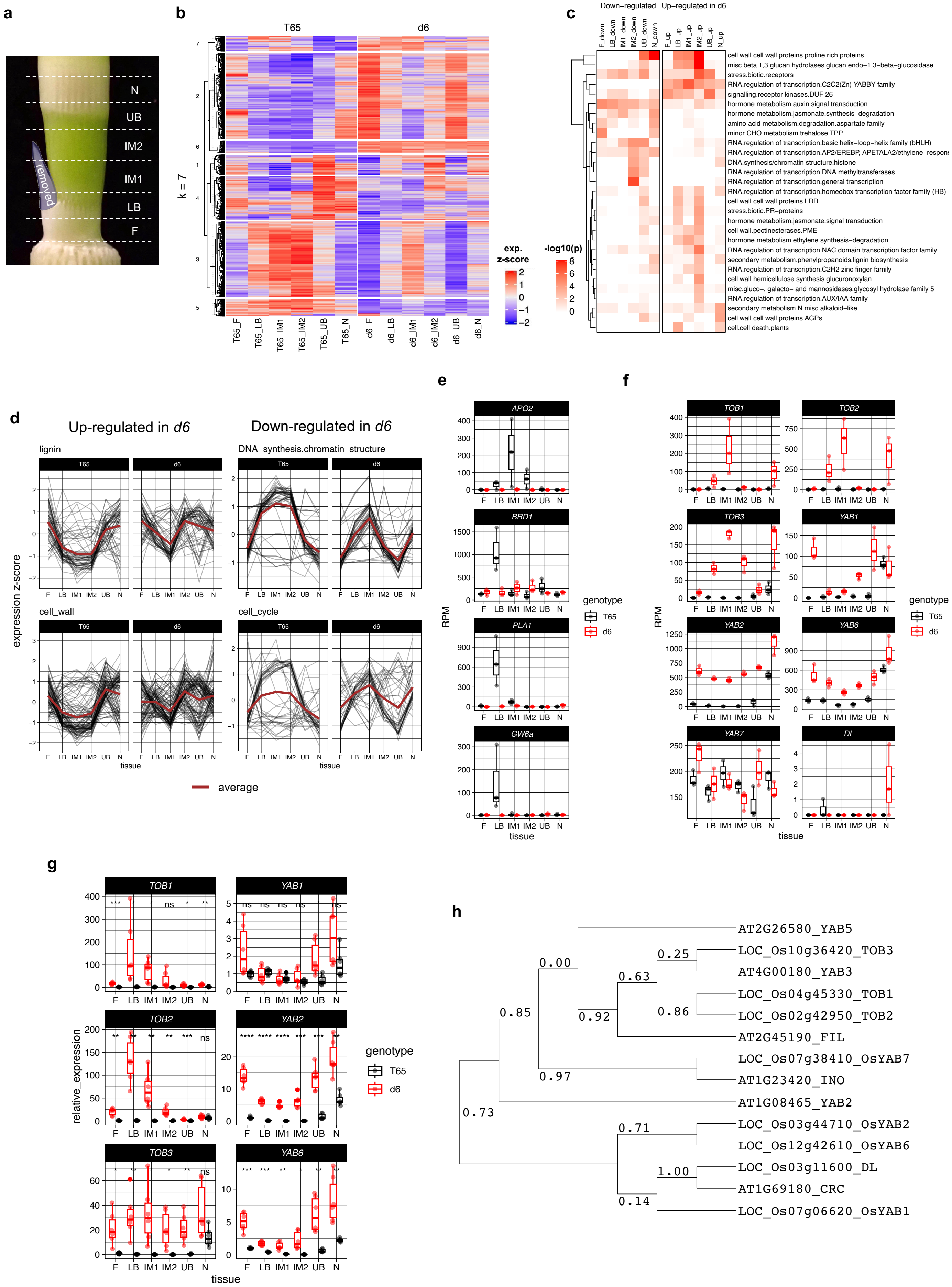
a
b
c
d
e
f
g
h
N
UB
IM2
IM1
LB
F
removed
T65
d6
k = 7
exp. z-score
-log10(p)
Down-regulated Up-regulated in d6
cell wall.cell wall proteins.proline rich proteins
misc.beta 1,3 glucan hydrolases.glucan endo-1,3-beta-glucosidase
stress.biotic.receptors
RNA.regulation of transcription.C2C2(Zn) YABBY family
signalling.receptor kinases.DUF 26
hormone metabolism.auxin.signal transduction
hormone metabolism.jasmonate.synthesis-degradation
amino acid metabolism.degradation.aspartate family
minor CHO metabolism.trehalose.TPP
RNA.regulation of transcription.basic helix-loop-helix family (bHLH)
RNA.regulation of transcription.AP2/EREBP, APETALA2/ethylene-respon
DNA.synthesis/chromatin structure.histone
RNA.regulation of transcription.DNA methyltransferases
RNA.regulation of transcription.general transcription
RNA.regulation of transcription.homeobox transcription factor family (HB)
cell wall.cell wall proteins.LRR
stress.biotic.PR-proteins
hormone metabolism.jasmonate.signal transduction
cell wall.pectinesterases.PME
hormone metabolism.ethylene.synthesis-degradation
RNA.regulation of transcription.NAC domain transcription factor family
secondary metabolism.phenylpropanoids.lignin biosynthesis
RNA.regulation of transcription.C2H2 zinc finger family
cell wall.hemicellulose synthesis.glucuronoxylan
misc.gluco-, galacto- and mannosidases.glycosyl hydrolase family 5
RNA.regulation of transcription.AUX/IAA family
secondary metabolism.N misc.alkaloid-like
cell wall.cell wall proteins.AGPs
cell.cell death.plants
Up-regulated in d6
Down-regulated in d6
lignin
DNA_synthesis.chromatin_structure
cell_wall
cell_cycle
expression z-score
tissue
average
APO2
BRD1
PLA1
GW6a
TOB1
TOB2
TOB3
YAB1
YAB2
YAB6
YAB7
DL
RPM
genotype
T65
d6
relative_expression
AT2G26580_YAB5
LOC_Os10g36420_TOB3
AT4G00180_YAB3
LOC_Os04g45330_TOB1
LOC_Os02g42950_TOB2
AT2G45190_FIL
LOC_Os07g38410_OsYAB7
AT1G23420_INO
AT1G08465_YAB2
LOC_Os03g44710_OsYAB2
LOC_Os12g42610_OsYAB6
LOC_Os03g11600_DL
AT1G69180_CRC
LOC_Os07g06620_OsYAB1

**Supplemental Figure 2. Transcriptome analyses in *d6* mutants.**

**a**, The schematic representation of stem samples. Each slice (1~2 mm) was dissected using a laser blade. F, foot II; LB, lower boundary; IM1, bottom part of the intercalary meristem; IM2, upper part of the intercalary meristem; UB, upper boundary; N, node II. **b**, Seven clusters of differentially expressed genes (DEGs, adjusted *p*-value < 0.01) based on the comparison between WT and *d6*. Genotypes and samples were shown at the top and bottom, respectively. The color range is shown in the expression z-score. **c**, Functional category enrichment analysis of DEGs. In the heatmap, DEGs down-regulated and up-regulated in *d6* mutants were shown on the left and right, respectively. The color range is shown in the $-\log_{10}$(*p-value*) (hypergeometric test). **d**, mRNAseq profiles of representative Functional categories. Brown lines are averages in each category **e**, mRNAseq profiles of known genes important for internode growth with three biological replicates. **f**, mRNAseq profiles of rice *YABBY* genes with three biological replicates. **g**, RT-qPCR validations of six *YABBY* genes with three biological and two technical replicates. ns, p>0.05; *, p<=0.05; **, p<=0.01; ***, p<=0.001; ****, p<=0.0001 (two-sided Student's t-test). In **e-g**, red and black transparent points, each sample; center line, median; box limits, upper and lower quartiles; whiskers, 1.5x interquartile range. **h**, A phylogenetic tree of Arabidopsis and rice YABBY proteins. Full-length amino acid sequences were aligned using mafft. The multiple alignment was trimed using Gblocks and the tree was inferred using PhyML inpremented in SeaView. Branch support was calculated with the aLRT option.

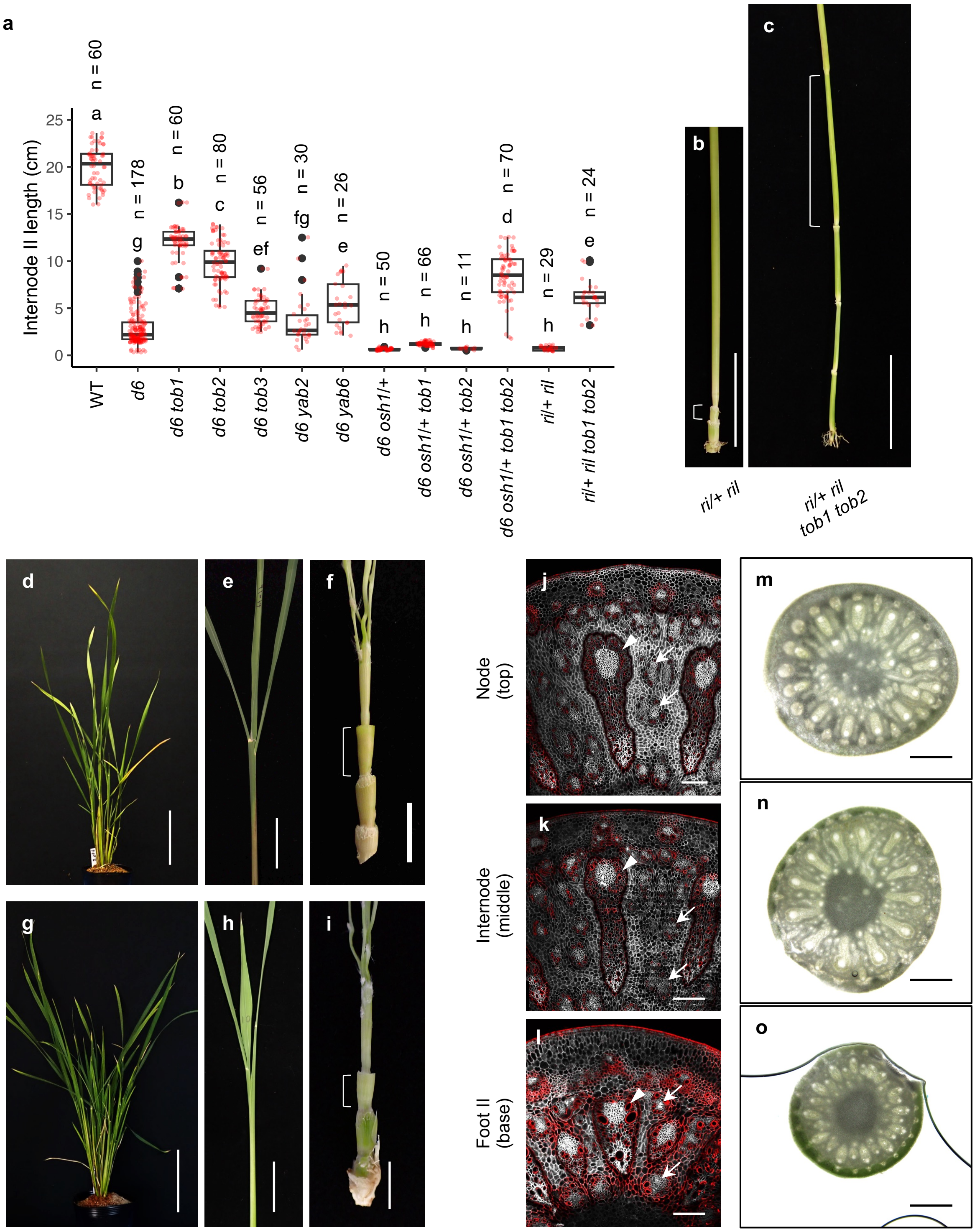


**Supplemental Figure 3. Stem phenotypes of mutants and *YABBY* overexpressors.**
**a**, Length of internode II in various mutant backgrounds. Different letters indicate significant differences ($p < 0.05$, Tukey-Kramer test). Red points, each sample; black points, outliers; center line, median; box limits, upper and lower quartiles; whiskers, 1.5x interquartile range. **b**,**c**, Stem phenotype in *ri/+ ril* mutants (**b**) and in *ri/+ ril tob1* and *tob2* mutations (**c**). **d-i,** Phenotypes of *TOB1* (**d-f**) and *TOB2* (**g-i**) overexpressors at maturity. In these plants, panicles failed to emerge from leaves (**e**, **h**), and stems were extremely short (**f**, **i**). Brackets indicate internode II. **j-l**, Transverse sections of *TOB1* overexpressor stems. Confocal images for basic fuchsin (red) and calcofluor white (gray) were merged. Arrows and arrowheads indicate ectopic diffusing vascular bundles and enlarged vascular bundles, respectively. **m-o**, Light microscopy images of transverse hand sections of *TOB2* overexpressor stems. Bars are 5 cm in **b**,**c**,**e**,**h**, 10 cm in **d**,**g**, 1 cm in **f**,**i**, 200 μm in **j-l**, and 500 μm in **m-o**.

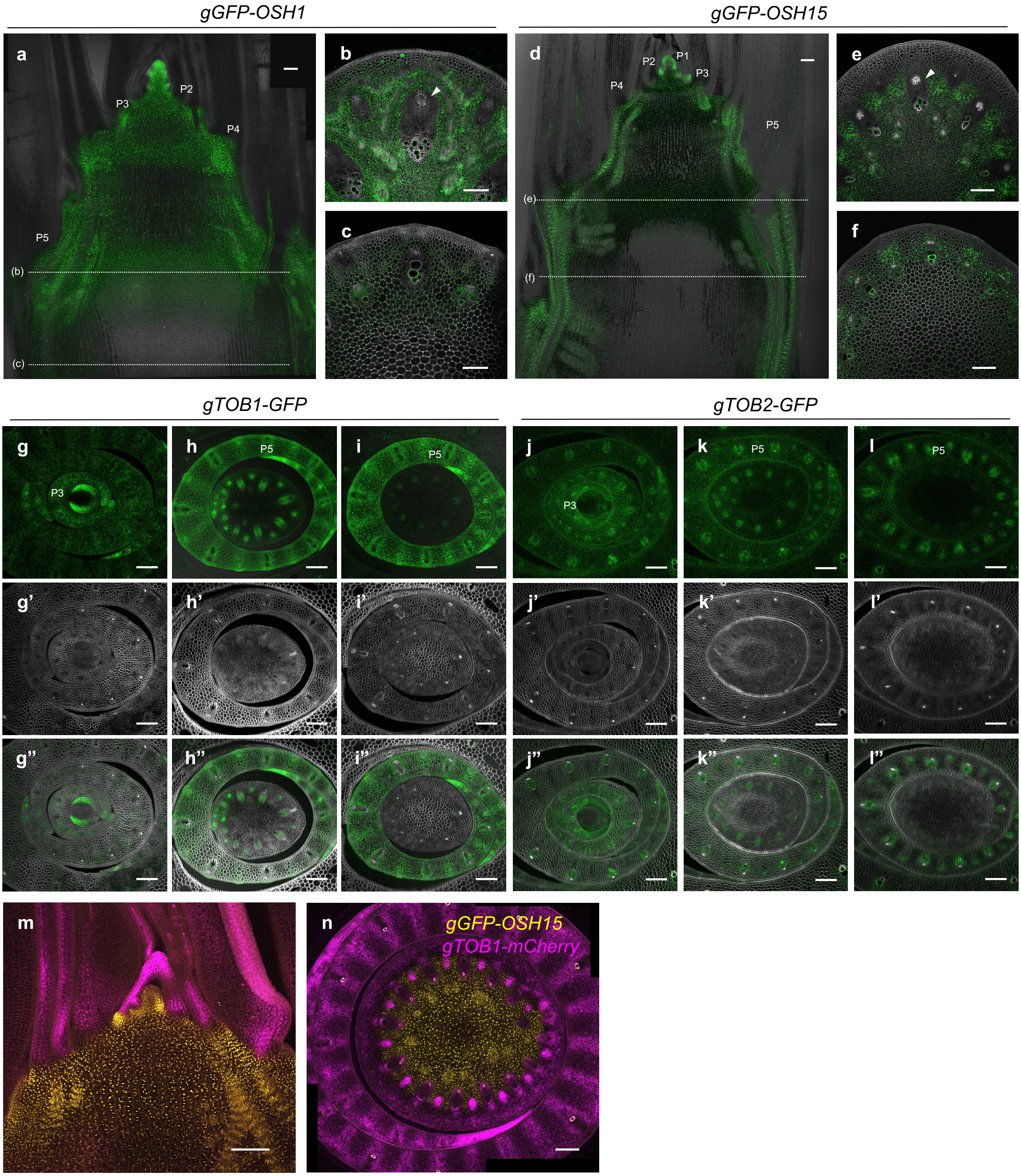


**Supplemental Figure 4. Expression of *KNOX1* and *YABBY* genomic reporters in the wild-type shoot apices.**

**a-f**, Confocal images of *gGFP-OSH1* (**a-c**) and *gGFP-OSH15* (**d-f**) reporters in the wild-type background. **a**,**d**, Longitudinal sections. The GFP channel (green) and blight field were merged. Dashed lines indicate approximate levels of transverse sections shown in **b**,**c**,**e**,**f**. **b**,**e**, Transverse stem sections at nodes. **c**,**f**, Transverse sections at internodes. Channels for GFP (green) and calcofluor white (gray) were merged. Arrowheads indicate enlarged vascular bundles. **g-l**, Transverse sections of *gTOB1-GFP* (**g-i**) and *gTOB2-GFP* (**j-l**) at the level of P2 node (**g**,**j**), P4 node (**h**,**k**), and P4 internode (**i**,**l**). **g-l**, **g'-l'** and **g''-l''** are channels for GFP (green), calcofluor white (gray), and merged images of them, respectively. **m**,**n**, Longitudinal (**m**) and transverse (**n**) sections of *gGFP-OSH15 gTOB1-mCherry* double reporters. Images of GFP (yellow) and mCherry (magenta) were merged. Bars are 100 μm.

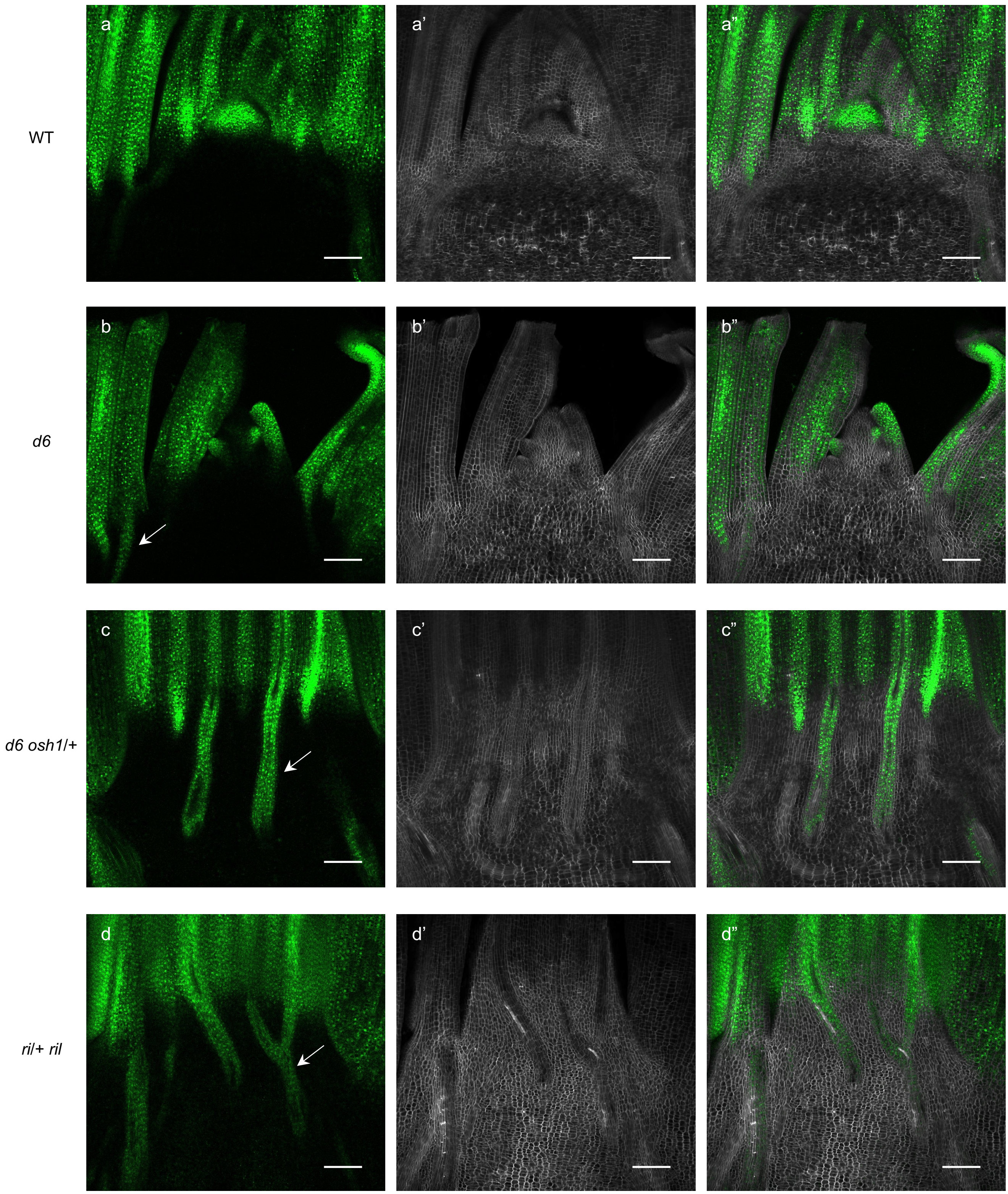


**Supplemental Figure 5. Expression of g*TOB1-GFP* in *knox1* and *blh* mutants.**
Longitudinal confocal images of *gTOB1-GFP* reporters in shoot apices around the reproductive transition. Backgrounds are wild type in **a**, *d6* in **b**, *d6 osh1*/+ in **c**, and *ri*/+ *ril* in **d**. Left (**a-d**), middle (**a'-d'**), and right (**a"-d"**) panels are GFP (green), calcofluor white (gray), and their merged images, respectively. Arrows represent ectopic expressions of *TOB1-GFP* along provascular bundles. Bars are 100 μm.

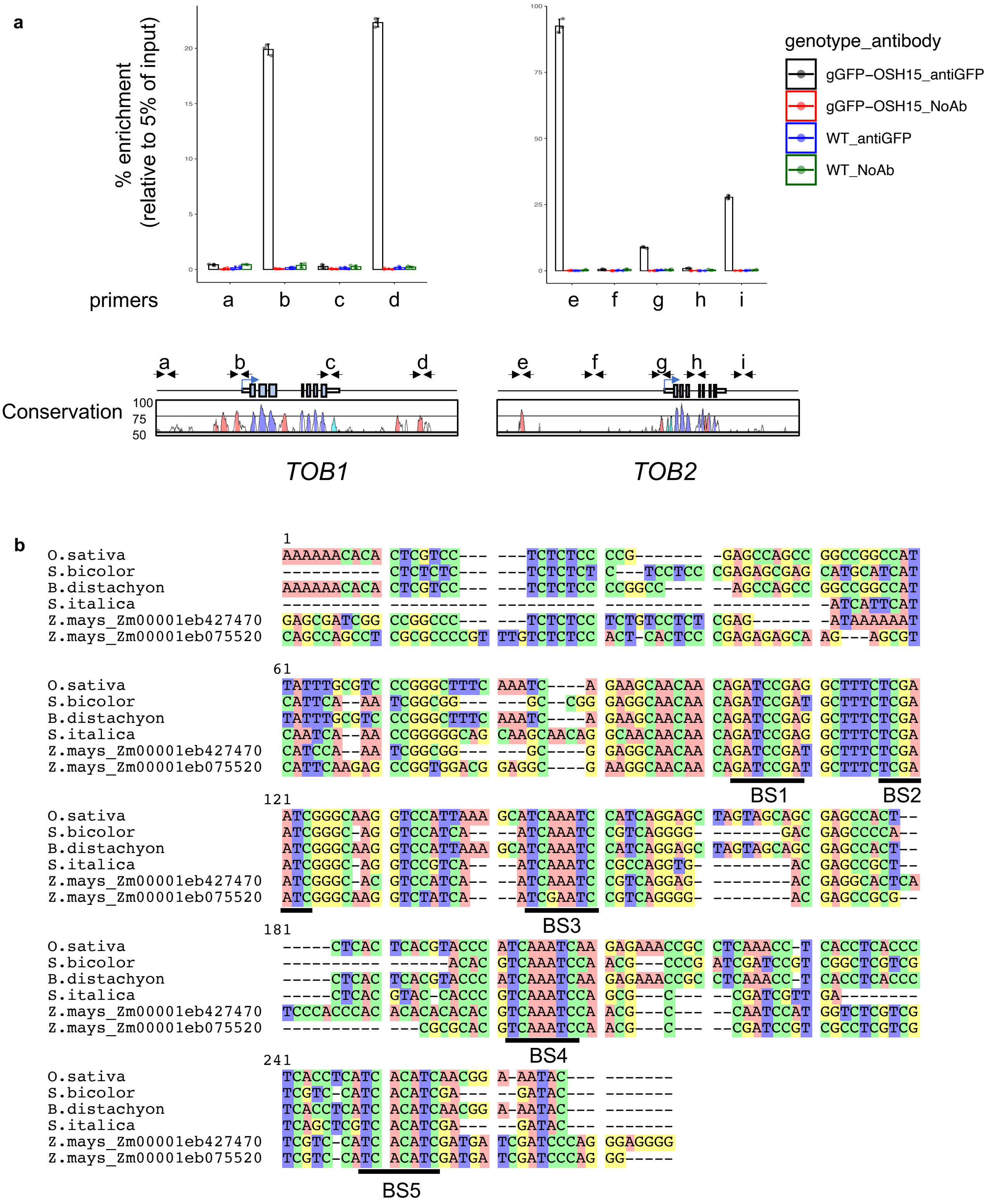


**Supplemental Figure 6. Validation of OSH15 binding to *TOB1/2* loci *in vivo* and nucleotide sequence alignment of the *TOB1* promoter.**
**a**, Validation by ChIP qPCR using gGFP-OSH15 young stems as samples and anti-GFP antibodies with three biological replicates. Dots, each sample; error bars, standard deviation. Primers used to detect enrichments of different *TOB1/2* regions were shown above gene models. **b**, Sequences of promoter regions from *TOB1* orthologs in other grasses were aligned with that from rice using Seaview software. Putative KNOX1 binding motif (GAnnnGA) were underlined.

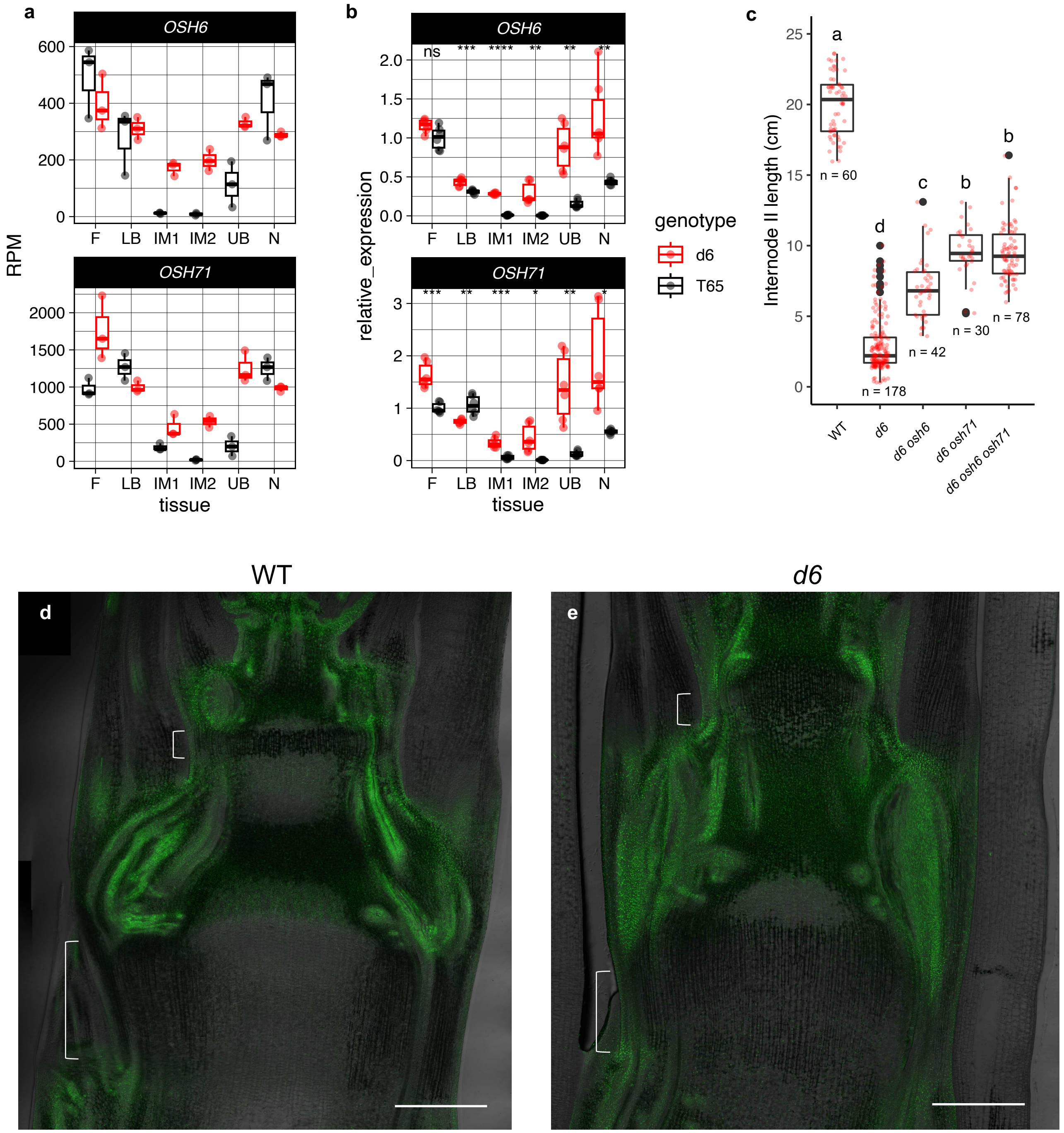


**Supplemental Figure 7. *OSH6* and *OSH71* were ectopically expressed in *d6* internodes and caused dwarfism.**

**a**, mRNAseq profiles of *OSH6* and *OSH71* with three biological replicates. **b**, RT-qPCR validations of *OSH6* and *OSH71* with three biological and two technical replicates. ns, p>0.05; *, p<=0.05; **, p<=0.01; ***, p<=0.001; ****, p<=0.0001 (Student's t-test). **c**, Length of mature internode II in *d6, osh6,* and *osh71* mutants. Numbers of stem samples were shown below the boxplots. Different letters indicate significant differences ($p < 0.05$, Tukey-Kramer test). **d,e,** Longitudinal confocal images of *gGFP-OSH6* reporters in the wild type (**d**) and *d6* mutants (**e**). Blight field- and GFP images were merged. Brackets indicate developing internodes. In **a-c**, red and black transparent points, each sample; black points, outliers; center line, median; box limits, upper and lower quartiles; whiskers, 1.5x interquartile range. Bars are 500 μm in **d**,**e**.

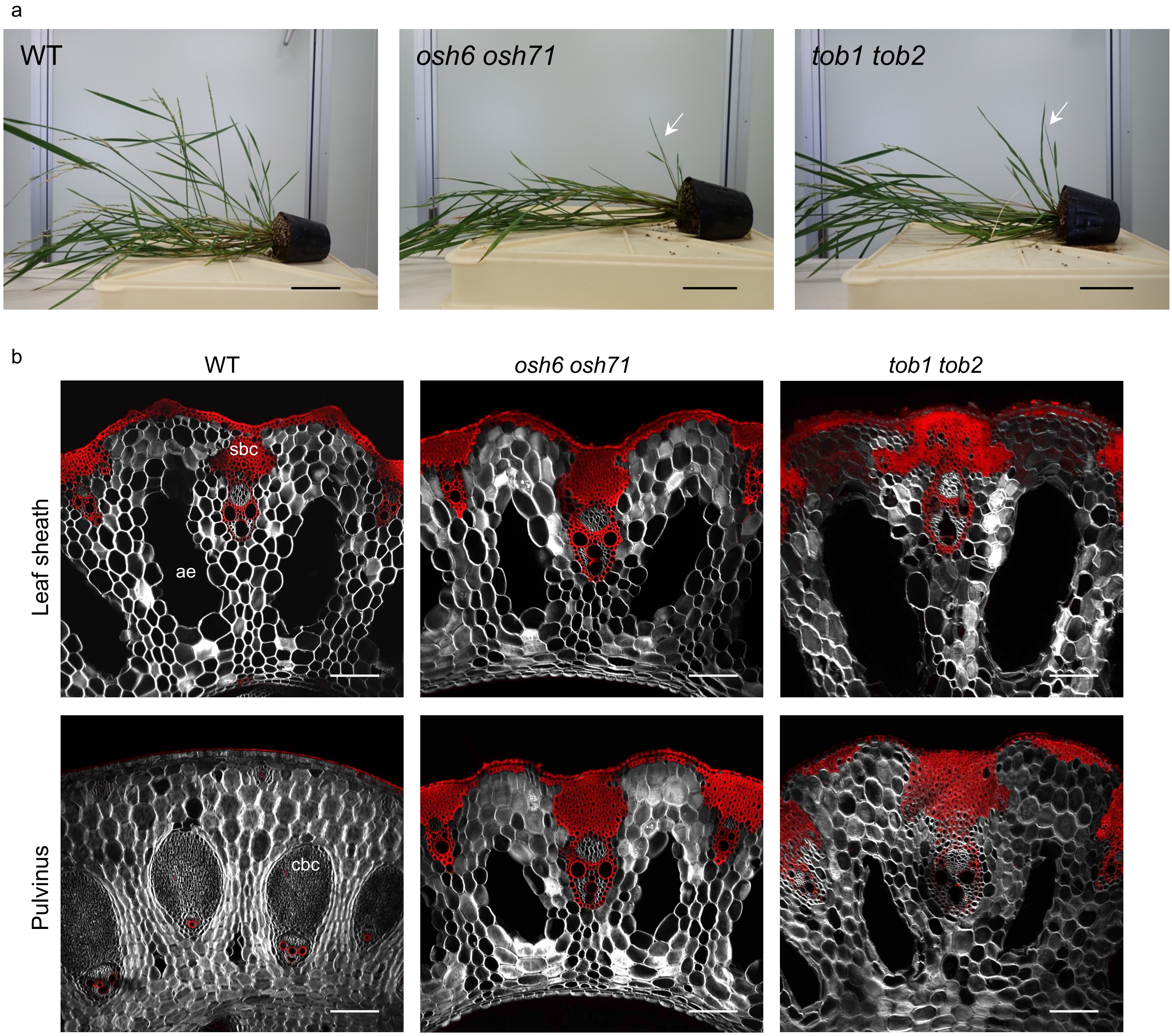


**Supplemental Figure 8. Pulvinus-less phenotypes and failure to respond gravity stimuli in *osh6 osh71* and *tob1 tob2* double mutants.**

**a**, Plant phenotypes 10 days after the onset of gravity stimuli. Arrows indicate newly formed tillers growing upright, indicating gravity sensing was not affected in these mutants. **b**, Transverse sections of mature leaves. sbc, sclerenchymatous bundle cap; ae, lysigenous aerenchyma; cbc, collenchymatous bundle cap. Confocal images for basic fuchsin (red) and calcofluor white (gray) were merged. Bars are 10 cm in a, and 100 μm in b.

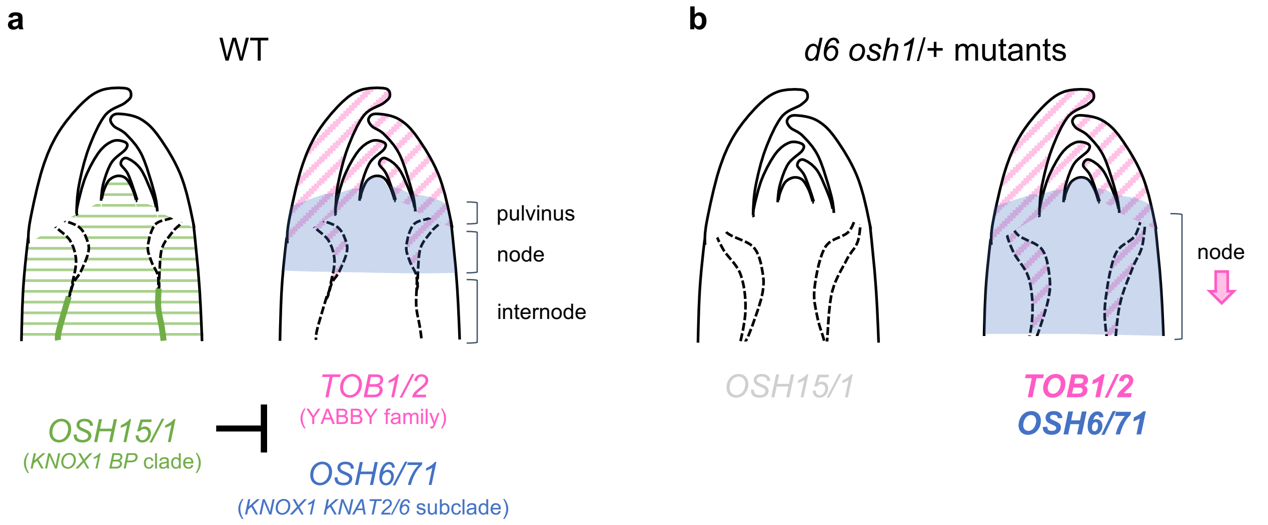


**Supplemental Figure 9. Schematic summary of *KNOX1* and *YABBY* functions in the control of the node-internode patterning.**

**a**, In the wild type, the *BP* subclade genes (*OSH15* and *OSH1*) repress *YABBY* genes (*TOB1* and *TOB2*) and *KNAT2/6* subclade genes (*OSH6* and *OSH71*) to confine the region of future nodes and thereby allow internode formation. **b**, In *d6 osh1*/+ mutants, *TOB1, TOB2, OSH6, and OSH71* expressions expand to the proximal direction and this results in the stem in which the node occupies the entire region. *YABBY* and *KNAT2/6* gene expressions overlap at the leaf base and induce pulvinus formation. Thus, *KNOX1* and *YABBY* gene expression patterns specify the subdomains of the stem.

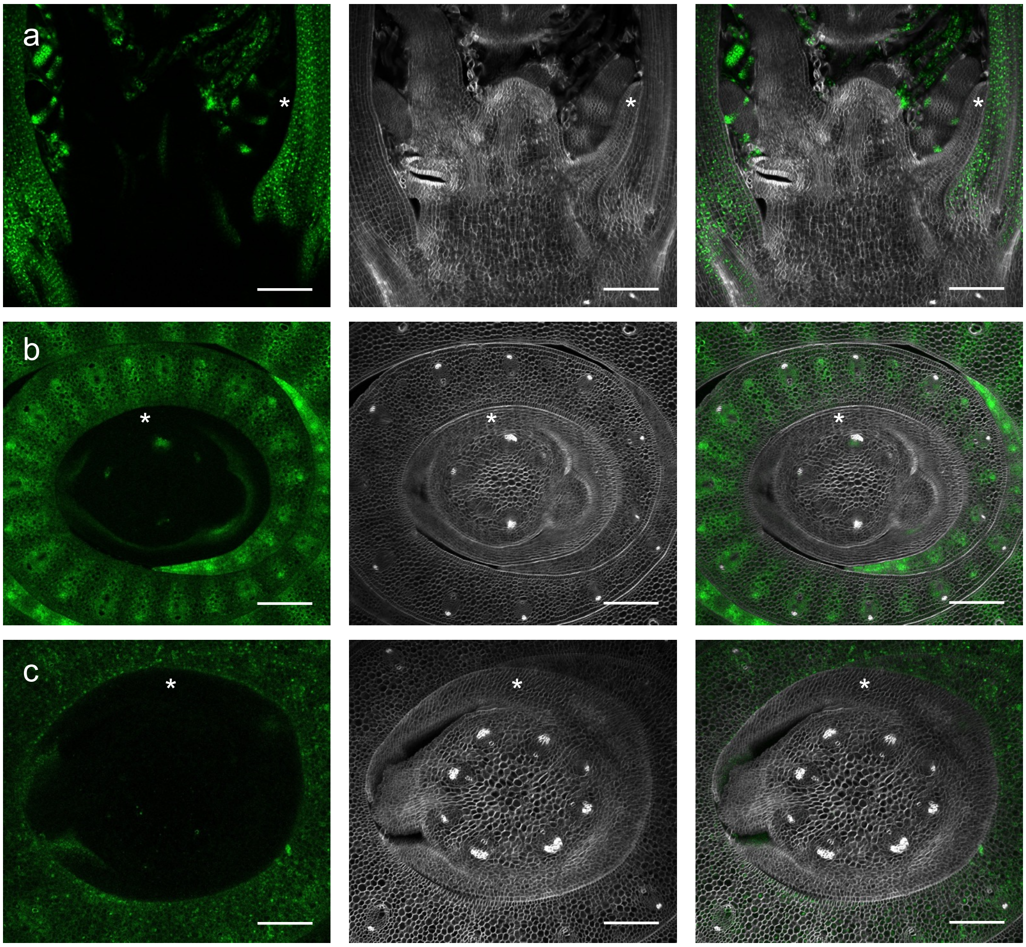


**Supplemental Figure 10. Expression of *TOB1* and *TOB2* in the inflorescence bract.**
**a,b**, *gTOB1-GFP* expression during early inflorescence development. **a** and **b** are longitudinal and transverse sections, respectively. **c**, *gTOB2-GFP* expression at a similar stage in a transverse section. The left and middle columns are GFP (green) and calcofluor white (grey) channels, and their merged images are on the right. Asterisks indicate bract primordia. Bars are 100 µm.
